# ChIP-seq of plasma cell-free nucleosomes identifies cell-of-origin gene expression programs

**DOI:** 10.1101/638643

**Authors:** Ronen Sadeh, Israa Sharkia, Gavriel Fialkoff, Ayelet Rahat, Jenia Gutin, Alon Chappleboim, Mor Nitzan, Ilana Fox-Fisher, Daniel Neiman, Guy Meler, Zahala Kamari, Dayana Yaish, Tamar Peretz, Ayala Hubert, Jonatan E Cohen, Salach Azzam, Mark Temper, Albert Grinshpun, Myriam Maoz, Samir Abu-Gazala, Ami Ben Ya’acov, Eyal Shteyer, Rifaat Safadi, Tommy Kaplan, Ruth Shemer, David Planer, Eithan Galun, Benjamin Glaser, Aviad Zick, Yuval Dor, Nir Friedman

## Abstract

Genomic DNA is packed by histone proteins that carry a multitude of post-translational modifications that reflect cellular transcriptional state. Cell-free DNA (cfDNA) is derived from fragmented chromatin in dying cells, and as such it retains the histones markings present in the cells of origin. Here, we pioneer chromatin immunoprecipitation followed by sequencing of cell-free nucleosomes (cfChIP-seq) carrying active chromatin marks. Our results show that cfChIP-seq provides multidimensional epigenetic information that recapitulates the epigenetic and transcriptional landscape in the cells of origin. We applied cfChIP-seq to 268 samples including samples from patients with heart and liver pathologies, and 135 samples from 56 metastatic CRC patients. We show that cfChIP-seq can detect pathology-related transcriptional changes at the site of the disease, beyond the information on tissue of origin. In CRC patients we detect clinically-relevant, and patient-specific information, including transcriptionally active HER2 amplifications. cfChIP-seq provides genome-wide information and requires low sequencing depth. Altogether, we establish cell-free chromatin immunoprecipitation as an exciting modality with potential for diagnosis and interrogation of physiological and pathological processes using a simple blood test.

**One Sentence Summary:** ChIP-seq of plasma-circulating nucleosomes (cfChIP-seq) from a simple blood test provides detailed information about gene expression programs in human organs, and cancer.

## Main Text

Genomic DNA is packaged into nucleosome complexes made up of ∼150bp DNA wrapped around histone proteins which are heavily post-translationally modified. The modifications are intimately coupled with transcriptional processes ^1–4^ --- monomethylation and trimethylation of Histone ^3^ Lysine ^4^ (H3K4me1 and, H3K4me3) mark active and paused enhancers and promoters respectively, and trimethylation of Histone ^3^ Lysine 36 (H3K36me3) marks elongation by RNA Pol II at gene bodies ^2,5–8^.

Upon cell death, the genome is fragmented and chromatin, mostly in the form of nucleosomes, is released into the circulation as cell-free nucleosomes (cf-nucleosomes) ^9–11^, that retain some histone modifications ^12–14^. We reasoned that capturing and DNA sequencing of modified nucleosomes from plasma may inform on a multitude of DNA-templated activities, including transcription, within the cells of origin (Figure 1A). This currently inaccessible epigenetic information extends beyond cfDNA modalities examined to date ^15–32^.

**Figure 1:**
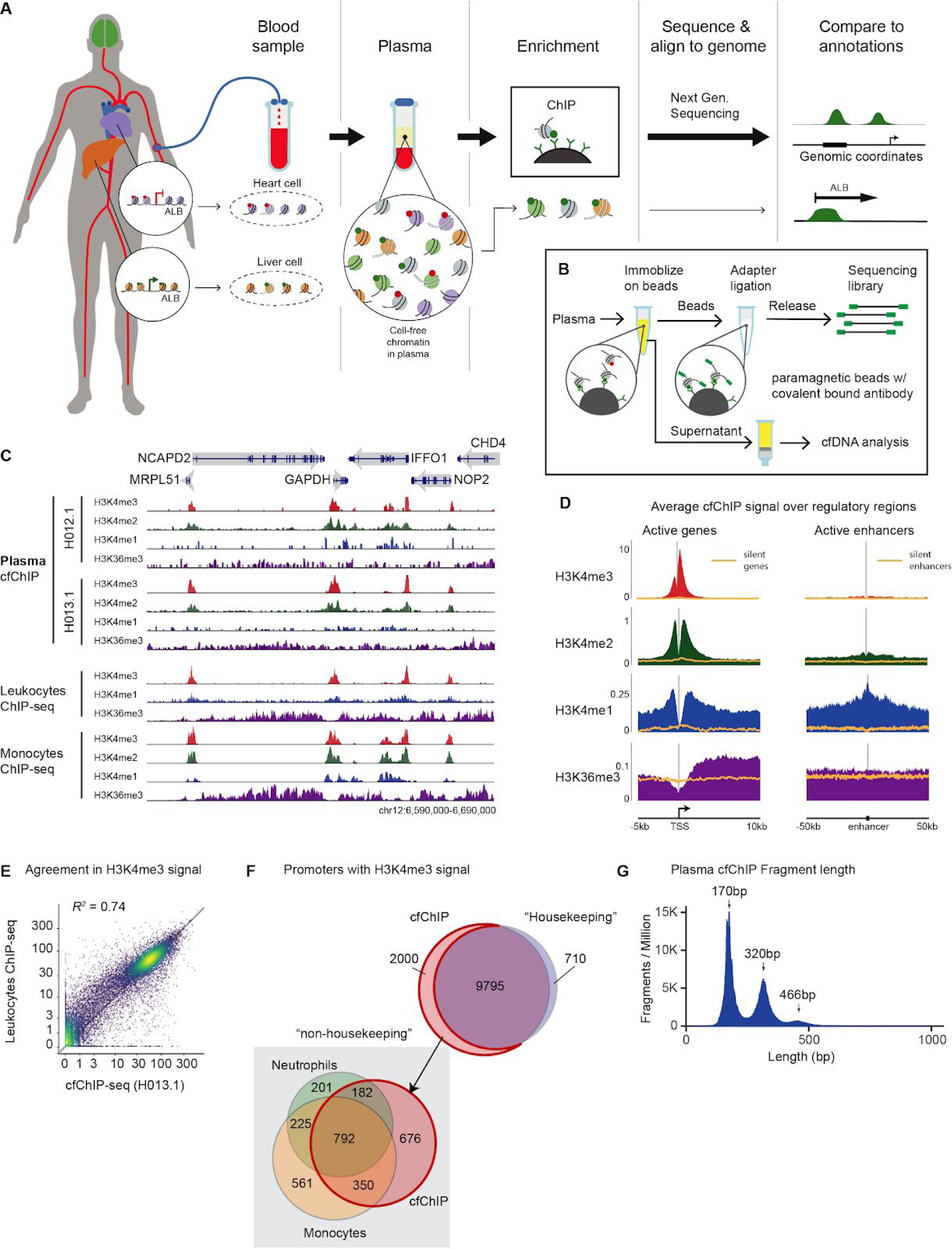
Immunoprecipitation of chromatin from plasma. A. cfChIP-seq method outline. Chromatin fragments from different cells are released to the bloodstream. These fragments are immunoprecipitated, and sequenced. Interpretation of the resulting sequences informs of gene activity programs in the tissue of origin. As an example, death of liver cells releases to the blood nucleosomes from the ALB promoter marked with H3K4me3. These are mixed with other circulating nucleosomes from other cells with the ALB promoter not marked by H3K4me3. After immunoprecipitation of H3K4me3 cf-nucleosomes and sequencing, we can detect fragments of DNA aligned to ALB promoters. These are indicative of death of hepatocytes since ALB promoter is marked by H3K4me3 only in hepatocytes. B. cfChIP-seq protocol. Antibodies are covalently bound to paramagnetic beads. Target fragments are immunoprecipitated directly from plasma. After washing, on-bead-ligation is performed to add indexed sequencing adapters to the fragments. The indexed fragments are released and amplified by PCR to generate sequencing-ready libraries. C. Genome browser view of cfChIP-seq signal on a segment of chromosome ^12^. Top tracks are cfChIP-seq signals from two healthy donors. The lower tracks are published ChIP-seq results from human white blood cells (leukocytes) ^4^. In each group we show four tracks corresponding to four histone marks -- H3K4me3 (red), H3K4me2 (green), H3K4me1 (blue), and H3K36me3 (purple). D. Meta analysis of cfChIP-seq signal over active promoters and enhancers. The orange line denotes the average of corresponding negative control regions (inactive genes and enhancers), providing an estimate of the background. Scale of all graphs is in coverage of fragments per million. E. Comparison of normalized H3K4me3 coverage of cfChIP-seq from a healthy donor against ChIP-seq from leukocytes ^4^. Each dot corresponds to a single gene. x-axis: healthy cfChIP-seq sample, y-axis leukocytes ChIP-seq. F. Analysis of promoters of RefSeq genes with a significant cfChIP-seq signal (methods) in healthy donors. cfChIP-seq captures most housekeeping promoters (ones that are marked in most samples in the reference compendium). The remaining 2000 non-housekeeping genes in cfChIP-seq show large overlaps with non-housekeeping promoters marked in neutrophils and monocytes, the two cell types that contribute most to cfDNA in healthy donors. G. Size distribution of sequenced cfChIP-seq fragments shows a clear pattern of mono- and di-nucleosome fragment sizes: x-ax1s: fragment length in base pairs (bp), y-axis: number of fragments per million in 1-bp bins.

Here, we perform Chromatin Immunoprecipitation and sequencing of cell-free nucleosomes directly from human plasma (cfChIP-seq). We show that cfChIP-seq recapitulate the original genomic distribution of marks associated with transcriptionally active promoters, enhancers, and gene bodies, demonstrating that plasma nucleosomes retain the epigenetic information of their cells of origin. We applied cfChIP-seq on ∼250 samples from more than hundred subjects including 61 self declared healthy donors, four patients with acute myocardial infarction, 29 patients suffering from autoimmune, metabolic, or viral liver diseases and 56 metastatic colorectal carcinoma (CRC) patients. We identified bone marrow megakaryocytes, but not erythroblasts, as major contributors to the cfDNA pool in healthy donors. We show pathology-related changes in hepatocytes transcriptional programs beyond changes in cells of origin. In CRC patients we detect the disease with high sensitivity and demonstrate that cfChIP-seq can identify subgroups of CRC patients with distinct cancer-related transcriptional programs, and with potential implications to diagnosis and treatment.

## Results

### ChIP-seq of cf-nucleosomes from plasma

We devised a simple protocol for cf-nucleosome ChIP-seq (cfChIP-seq) from small amounts of plasma --2ml of plasma from healthy donors and <0.5ml from patients with increased levels of cfDNA (Methods). Briefly, to overcome the extremely low concentration of cf-nucleosomes and the high concentration of native antibodies in plasma, we incorporated two modifications to standard ChIP-seq protocols (Figure 1B). First, we covalently immobilized the ChIP antibodies to paramagnetic beads, which can be incubated directly in plasma avoiding competition with native antibodies. Second, we maximize efficiency by using an on bead adaptor ligation ^33–36^, where barcoded sequencing-DNA adaptors are ligated directly to chromatin fragments prior to the isolation of DNA. The resulting protocol allows us to simply and efficiently enrich and sequence targeted chromatin fragments from low volumes of plasma.

We performed cfChIP-seq on multiple plasma samples from healthy individuals with antibodies targeting marks of accessible/active promoters (H3K4me3 or H3K4me2), enhancers (H3K4me2, or H3K4me1), and gene body of actively transcribed genes (H3K36me3) (Figure 1C). cfChIP-seq profiles obtained from on the same blood sample with different antibodies show the expected patterns (Figures 1C, 1D). While we see H3K4me3 cfChIP-seq signal almost exclusively at promoters, a large fraction of the reads from H3K4me2 cfChIP-seq are mapped to putative enhancer regions (Methods, Figure S1A).

cfChIP-seq is highly specific, as argued by several lines of evidence: (a) cfChIP-seq signal is consistent with reference ChIP-seq in tissues ^4^, evident by the remarkable agreement of peaks in genome browser (Figures 1C, S1B), in the average pattern around promoters and enhancers (Figures 1D, S1C), and in quantitative comparison of the signal across multiple genomic locations, such as all promoters, (*R* > 0.8 Figures 1E, S1D). Essentially all promoters that are ubiquitously marked (housekeeping) by H3K4me3 in reference ChIP-seq are significantly enriched for this mark in cfChIP-seq (9,795/10,505 promoters 93%, *p* < 10-1000). Focusing on marked promoters from non-housekeeping genes in cfChIP-seq, there is significant overlap (1,324/2,311 promoters 57%, *p* <10-288) with promoters from monocytes and neutrophils that are the major contributors to the cfDNA pool ^16,31^ (Figure 1F). (b) Performing cfChIP-seq with a mock antibody resulted in dramatically lower yield, without the enrichment seen for histone modifications (Table S1). (c) We estimated the rate of non-specific events in each sample (Methods) and used this background noise model to evaluate the expected amount of signal originating from a non-specific source in each assay (Methods, Table S1). These results show that for H3K4me3 the levels of non-specific reads are comparable to or lower than reference ChIP (Figure S1E) while for other antibodies such as H3K36me3 the performance is reduced, but remains highly informative (below).

The signal obtained by cfChIP-seq is not due to white blood cells lysed during sample handling. Several avenues of evidence show this. (a) Fragment size distributions of cfChIP-seq correspond to DNA wrapped around mono- and di-nucleosomes (Figures 1G, and S2A), consistent with apoptotic or necrotic cell death, but not with cell lysis, which results in much larger (>10kb) fragments ^37^. (b) We identified 676 promoters carrying H3K4me3 that are absent in ChIP-seq from white blood cells (leukocytes, peripheral blood mononuclear cells; Figures 1F), these include promoters of genes that are expressed specifically in megakaryocytes, which reside in the bone marrow (below). (c) In patients, we detect disease-related chromatin from remote tissues including heart, liver, and colon (below).

Together, these results strongly suggest that cf-nucleosomes originate in cells that have died *in vivo,* and preserve the endogenous patterns of active histone methylation marks in the cells of origin and can be assayed by cfChIP-seq.

### cfChIP-seq detects changes in cell free nucleosome origins

To appreciate the variability in cfChIP-seq signals between samples, we examined several self-reported healthy donors. We found a high similarity of signal between individuals that was in the range of the observed similarity between samples obtained on different days from the same donor (Figure S2B, Supplemental Note). We next turned to examine the differences between samples from healthy donors and samples from a cancer patient with advanced stage metastatic CRC. We expected that a large fraction of the cfDNA in advanced cancer patients will be from tumor origin ^18,31^. Indeed, samples from this patient had a substantial increase in cfDNA concentration (84-122 ng/ml vs 4-10 ng/ml for healthy donors), the majority of which is most likely of tumor, or tumor-adjacent origin.

Comparing these cancer samples to healthy samples we observe dramatic differences in cfChIP-seq signal of H3K4me3, H3K4me2, and H3K36me3 (Figure 2A). These include specific, and statistically significant increases in H3K4me3 (1,639 regions), H3K4me2 (2,595 regions), and H3K36me3 (5,416 regions) (Methods). Many of these regions show no signal in samples from healthy donors. Genes associated with these regions include several classic CRC markers, such the long non-coding RNA CCAT1 (colorectal cancer associated transcript 1)^38^, CRC-specific transcription factors such as CDX1, and the carcinoma associated antigen EPCAM (Figure 2A). In addition, we observed increased H3K4me3 modification at the promoter of the non-coding antisense RNA EGFR-AS1 ^39^. While H3K4me3 cfChIP-seq signal at the promoter for EGFR is detected in both healthy and cancer samples, EGFR-AS1 is detected only in the cancer patient. This finding, which can not be detected by cfDNA mutation analysis, highlights the potential relevance of cfChIP-seq for treatment choice beyond genomic mutations.

**Figure 2:**
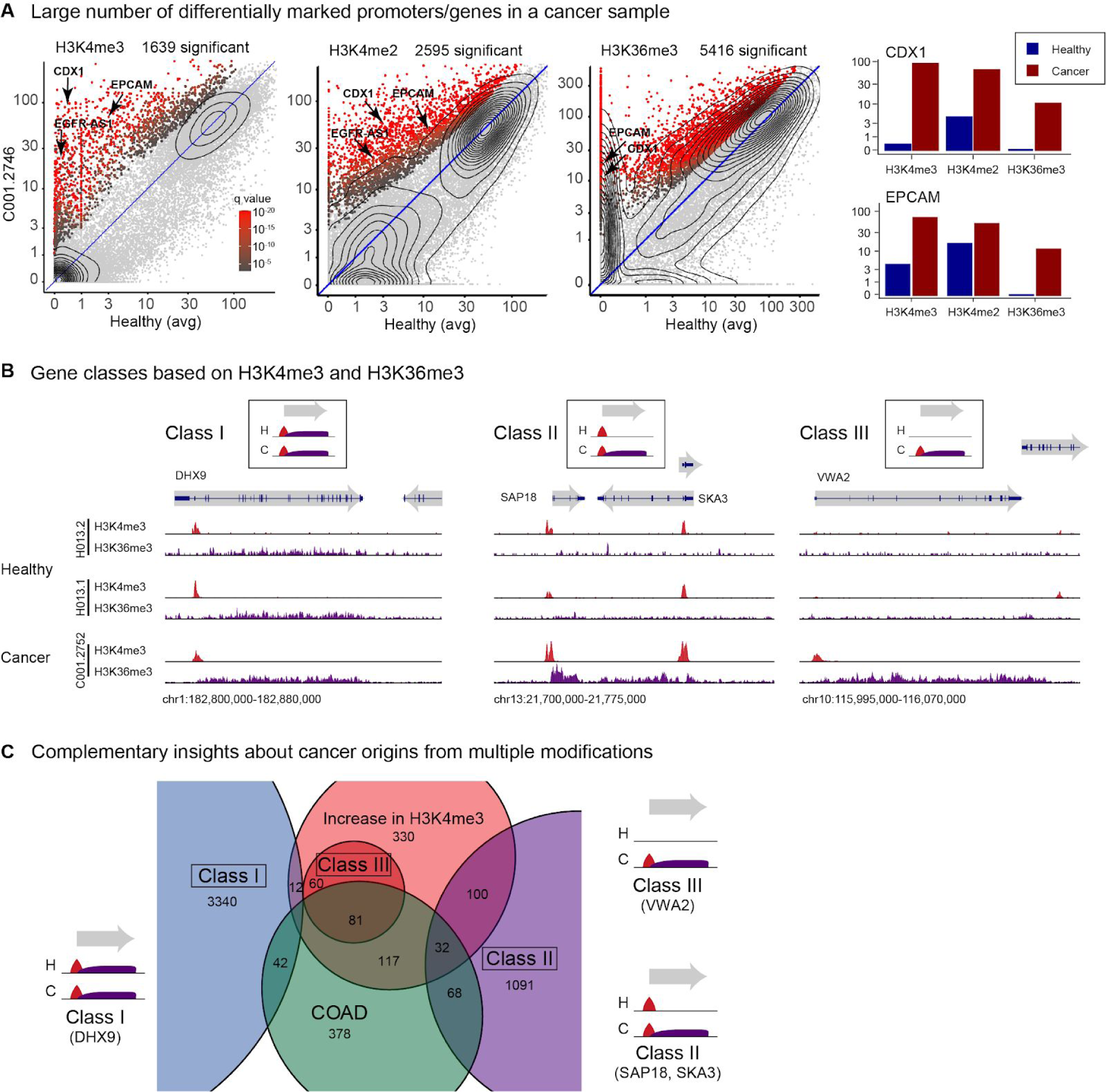
cfChIP-seq of multiple marks is informative on gene expression. A. Detection of genes with significant high coverage in a sample from a colorectal cancer (CRC) patient (Table S3). For each gene we compare mean normalized coverage in a reference healthy cohort (x-axis) against the normalized coverage in the cancer sample (y-axis). For H3K36me3, the signal is normalized by gene length. Significance test whether the observed number of reads is significantly higher than expected based on the distribution of values in healthy samples (Methods). Example of the levels of two genes in these comparisons are shown on the bar chart (right panel). B. Browser views of genes that demonstrate different H3K4me3 and H3K36me3 classes. Class I: genes marked by both marks in healthy and cancer patient samples. Class II: genes marked by H3K4me3 in healthy and cancer samples, but with H3K36me3 only in the cancer patient sample (gain of H3K36me3). Class III: genes marked by both marks only in the cancer patient sample (gain of both marks). C. Venn diagram (zoom in view) showing the relations of genes from the three classes in B with the set of genes that show increased H3K4me3 and the set of genes previously identified to be highly expressed in colorectal adenocarcinoma (COAD, Methods).

cfChIP-seq signals are enriched with promoters and genes related to CRC and not other cancer types. To systematically test for cancer-specific signatures in cfChIP-seq signal we analyzed expression profiles from The Cancer Genome Atlas (TCGA) and GTEx projects ^40,41^. For each cancer type we generated a cancer-specific signature composed of the set of genes whose expression is significantly higher in the tumor compared to normal tissues (Methods, Table S2). We then tested for significant overlaps between the set of genes with a higher cancer-specific cfChIP-seq signal and the set of genes in each cancer signature (Methods). The set of genes with high H3K4me3 signal in the cancer patient has a significant overlap (303 of 739 genes, *q* < 10-90) with GI-tract adenocarcinoma genes (COAD), but only a negligible overlap with non-GI cancers such as diffuse large b-cell lymphoma (DLBC) (Figure S3A).

Tissue-specific enhancers can also be detected by cfChIP-seq. Since H3K4me2 marks promoter proximal and enhancer regions we examined how H3K4me2 signal at enhancers differentiates the cancer sample from samples of healthy donors. To ensure that we are not biased by promoter signal, we focused on regions of a size larger than 600bp that are at least 5Kb from the nearest TSS and do not overlap a gene body. This smaller set of regions (48,525/2,345,831 regions) can be safely assumed to be of enhancers. Using the Roadmap Epigenomics compendium chromatin annotations, we assigned for each cell-type a set of unique distal enhancers. Comparing H3K4me2 signal in healthy samples to a sample from a colorectal cancer patient, we observed significantly higher signal in colon-specific enhancers, which are barely present in healthy samples (Figure S3B).

The activity of elongating RNA polymerase at gene bodies can be monitored by H3K36me3 cfChIP-seq. Unlike H3K4me3, which marks transcription start sites at both poised and active genes, Tri-methylation of H3 lysine ^36^ (H3K36me3) requires active transcription elongation to be deposited, and is hence more indicative of gene activity ^2^. Despite the high background of H3K36me3 in cfChIP-seq (Figure S1D), we do observe the typical enrichment at gene bodies (Figures 1D and S3C) and the signal in healthy donors correlates with leukocyte RNA-seq (Figure S3D). Comparing the H3K36me3 signal from a healthy donor to that of a colorectal adenocarcinoma patient, we observe 5,416 genes that are hyper H3K36 tri-methylated, by at least ^4^ fold in the cancer sample compared to healthy donors (Figure 2A).

The signal from H3K4me3 and H3K36me3 cfChIP-seq corroborates the cancer origin of nucleosomes. Examining the ∼5400 genes with increased H3K36me3 signal in this cancer sample, we distinguish between three main classes. Class I includes ∼3,400 genes that are marked by both H3K36me3 and H3K4me3 in healthy and cancer samples (e.g., DHX9, Figure 2B). Class II contains ∼1,300 genes that are marked with H3K4me3 in both healthy and cancer samples (e.g., SAP18 and SKA1, Figure 2B) but differ in H3K36me3 signal, which provides new information beyond H3K4me3. Finally, 159 Class III genes are not marked with either signal in healthy samples (e.g., VWA2, Figure 2B). Contrasting the set of highly expressed COAD signature genes, with these three classes, we observe that each class captures different parts of these sets (Figure 2C). Specifically, ^68^ COAD genes that were not differentially marked by H3K4me3 are detected as active by H3K36me3. Moreover, for 113 COAD genes (32 in Class II and ^81^ in Class III) the change in H3K4me3 signal is further corroborated by H3K36me3 signal.

Altogether, these results demonstrate the ability of cfChIP-seq to probe the state of various genomic features including promoters, enhancers, and gene bodies. Moreover cfChIP-seq detects functional changes of these features in samples from a cancer patient. The observed changes are consistent with independent studies of this type of cancer. These results strongly support the notion that the nucleosomes captured by, cfChIP-seq were modified in, and originate from the remote solid tissue. The multiple modification assays and their analysis demonstrates that each of these features is highly informative on various aspects of transcriptional activity in the cells of origin.

### cfChIP-seq is highly sensitive

The ability of cfChIP-seq to detect rare molecular events in the cfDNA pool is dictated by several factors: 1. The number of informative molecules in the sampled plasma; 2. The capture rate of marked informative molecules; and, 3. The signal to noise ratio (SNR) of the assay. We examine each of these in detail.

The number of informative molecules in the plasma depends on the number of the relevant marked nucleosomes in each cell of interest. This number is proportional to the size of the genomic region in question and the amount of cells of interest that had shed their nucleosomes to the blood (Figure 3A). For example, there are ^30^ cardiomyocyte-specific promoters with varying lengths that consist of 366 nucleosomes that are marked with H3K4me3 only in cardiomyocytes, these promoters drive expression of many cardiomyocyte-specific genes such as the troponins TNNT2 and TNNI3 and the cardiac myosin heavy chain MYH6. All the marked molecules originating from these regions in cardiomyocytes are informative for detecting cardiomyocyte presence. Assuming a 1% contribution of cardiomyocyte to a cf-nucleosomes pool of ∼1,000 genomes/ml, we expect ∼7,320 informative nucleosomes in a 2ml sample. Estimates for different cell types are given in (Figure S4A).

**Figure 3:**
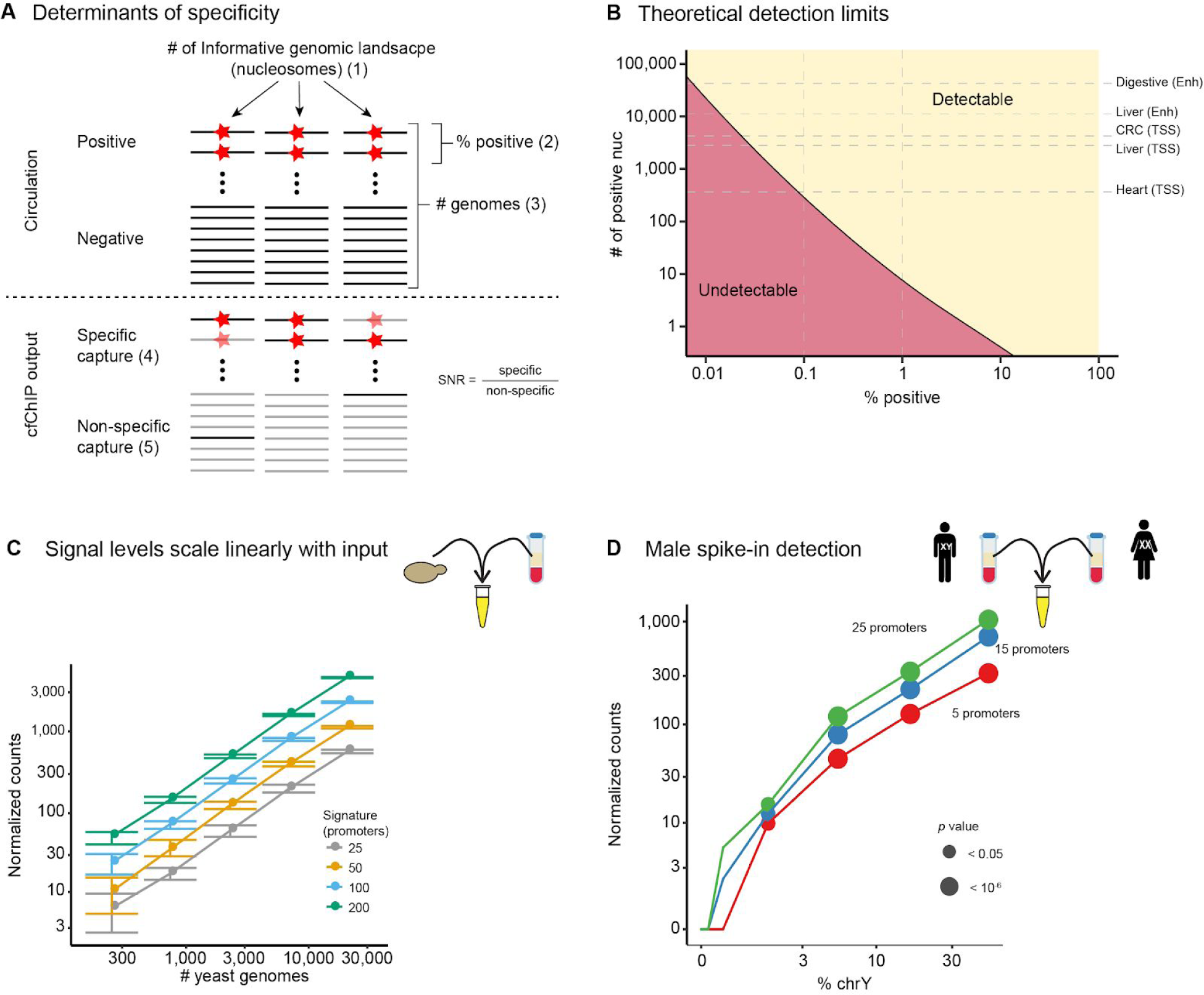
cfChIP-seq sensitivity. A. Schematics of the parameters involved in determining cfChIP-seq sensitivity. 1. **Number of informative nucleosomes** is the total number of signature-specific nucleosomes in the plasma that carry a mark of interest; 2. The **percent** of fragments contributed by the signature-positive cells among the fragments in circulation; 3. Total **number of genomes** in circulation; 4. The **specific capture probability** of marked nucleosomes by the cfChIP-seq assay; and 5. The **non-specific capture probability** of nucleosomes (background). The signal to noise ratio (SNR) is the ratio of the specific to non-specific capture probabilities. B. Simulation analysis of event detection power as a function of percent positive (x-axis) and number of informative locations (y-axis). Detection is defined as 95% probability of assay results (capture & sequencing) that reject the null hypothesis of background signal with p < 0.05 (Methods). Simulation assumes #number of genomes = 10,000 (10 ml plasma of healthy donor), capture probability of 1%, and SNR of 500. The size of several example signatures are shown (see text). C. Signal level is linear with input. Plasma of a healthy donor was spiked in with different amounts of yeast nucleosomes (x-axis). The number of counts observed (y-axis) for signatures of different sizes. Error bars show 20-80% range over 100 different sampled signatures of the given size. D. Test of sensitivity using male spike-in. Plasma of healthy female and male donors were titrated at different ratios. Detection of male-specific promoters as a function of percent of chrY genomes in the sample (x-axis). Shown are the number of counts (y-axis) and significance (circle radius).

To evaluate the capture rate of marked molecules, we used our prior knowledge of the genomic distribution of H3K4me3 marked nucleosomes, which are highly localized to transcription start sites (Figure 1C, D). Using this prior knowledge we distinguish between non-specific capture (in regions without TSSs) to specific capture (in TSSs that are known to be constitutively marked by H3K4me3). We can then contrast the amount of input molecules against the number of uniquely sequenced reads to estimate the probability of specific capture and the probability of nonspecific capture (Figures S4B and S4C, Methods). With these assumptions and by using two different approaches (Methods), we estimate the H3K4me3 specific capture probability to range between 0.01% and 0.1% across dozens of cfChIP-seq experiments (Figure S4D).

Based on these estimates we modeled the detection probability for a signature as a function of the percent contribution to the cf-nucleosome pool, and the number of informative nucleosomes (Figure 3B). This estimate assumes independence of the concentration of plasma nucleosomes and capture rate. This assumption was verified by spiking increasing amounts of yeast-derived nucleosomes into a healthy donor plasma (Figure 3C). Assuming 10ml of plasma, which is standard in the field, and using realistic background and capture rates, we predict that detection of a cell-type that contributes 0.1% to the cf-nucleosome pool, requires >200 uniquely marked nucleosomes (>50 promoters). This is much lower than the number of informative H3K4me3 nucleosomes in several tissue signatures such as heart, liver, or CRC, and an order of magnitude lower than the number of potential informative H3K4me2, and H3K36me3 nucleosomes (Figures 3B and S4A). Such a sensitivity approaches the range that is thought to be required for early detection of cancer via ctDNA ^42^.

To evaluate these predictions in practice, we took advantage of sequences unique to the Y chromosome and titrated male-derived plasma into female-derived plasma. We evaluated the sensitivity for genomic signatures of different sizes at male-specific locations on the Y chromosome (Figures 3D and S4D), detecting the presence of male chrY when it was present in 1.5% of the genomes in the plasma (∼30 copies, Figure 3D). These numbers are consistent with our estimates based on the parameters of the specific experiment (Figure S4E and S4F).

This collection of technical experiments, and their analysis, establish the quantitative nature of the method, provide a robust and cross-validated estimate of the sensitivity of the assay and the main parameters underlying this sensitivity. Importantly, this analysis assumed a single (and rare) modification is assayed, however, a combination of several modifications will no doubt increase the effective sensitivity, and bolster confidence in specific co-occurring findings, as we demonstrated (above)..

### cfChIP-seq of H3K4me3 correlates with gene expression

Having established that cfChIP-seq captures active histone modifications, and identifies differential modification states in diseased tissues, we decided to systematically evaluate the extent to which these reflect gene expression patterns in the cells of origin. We focused on the well characterized H3K4me3 mark since the signal is highly focused at promoters and is predictive of gene expression levels ^43–45^.

To validate the relationship between promoter H3K4me3 and gene expression levels, we used ^56^ Roadmap Epigenomics samples that have matching gene expression and H3K4me3 ChIP-seq profiles. For each gene, we compared the expression levels of the gene to promoter H3K4me3 ChIP-seq signal across all samples (Methods). We find that for a large group of genes (10,150/14,313 genes), H3K4me3 ChIP-seq signal is significantly correlated with differences in expression levels of the gene (pearson 0.28≤*r*≤0.99; Figure 4A). Most of the genes that do not have significant correlation are either genes that have high H3K4me3 levels in their promoters in most samples (housekeeping, 1,616/4,163 genes, e.g., RAD23A) or genes with low levels of expression in all tissues (1,299/4,163 genes).

**Figure 4:**
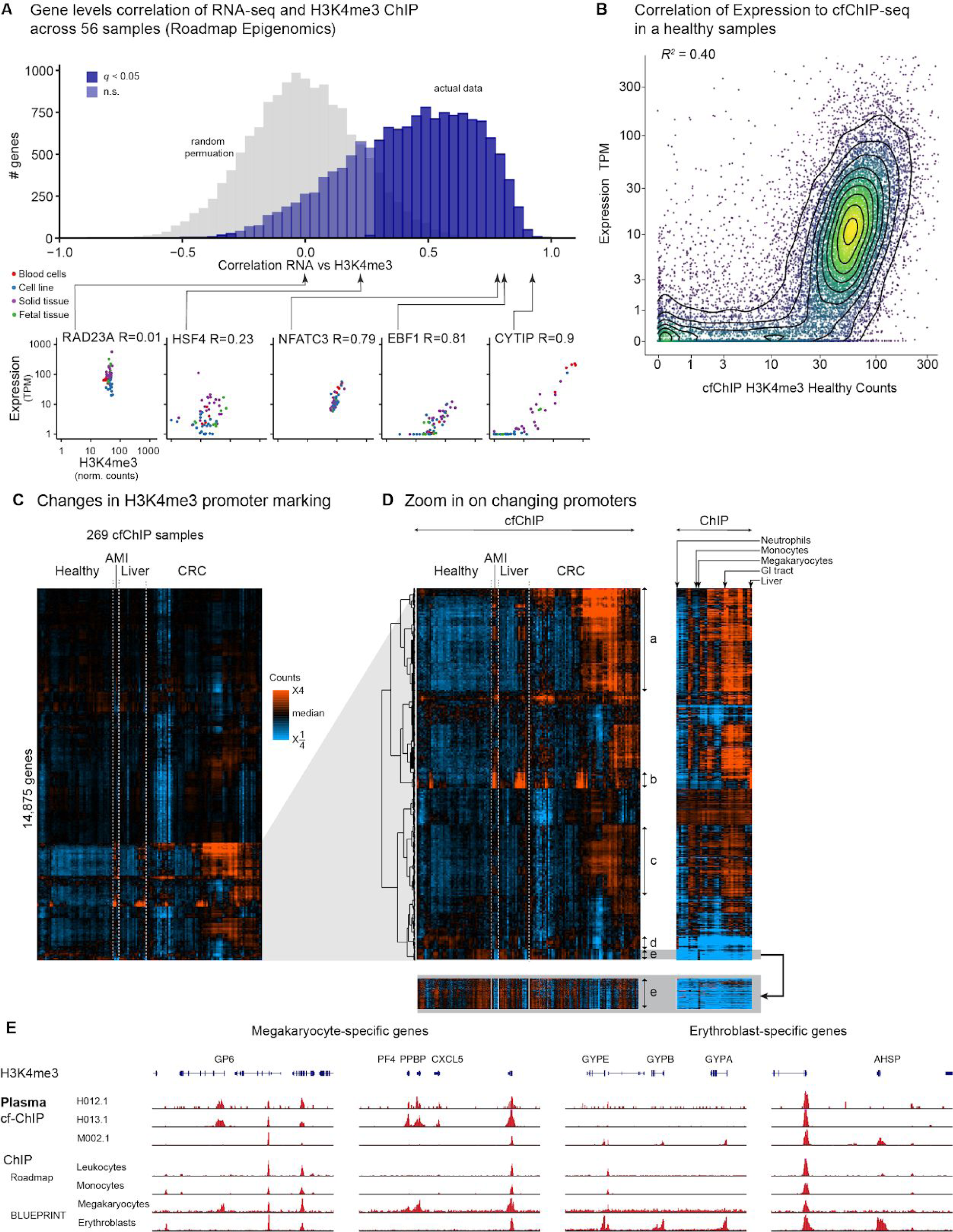
H3K4me3 cfChIP-seq signal is correlated with expression levels. A. Gene level analysis of the correlation in expression level and H3K4me3 methylation levels across ^56^ Roadmap Epigenomic samples ^4^ where we have matching profiles of both expression and H3K4me3. For each gene we computed the Pearson correlation of its normalized expression levels and normalized H3K4me3 levels (as computed by our pipeline) across the samples. Shown is a histogram of the correlations on all RefSeq genes. In gray we show the histogram resulting from random permutation of the relation between expression profiles and H3K4me3 profiles. Below: examples of genes with different correlation values. High correlation genes such as CYPIP, NFAT3, and EBF1 show coordinated change in both values. Low correlation genes either have little dynamic range in one of the measures (e.g., RAD23A is highly expressed in all tissues and has roughly constant H3K4me3 levels) or are not related (e.g., HSF4). The latter can be due to annotation errors of promoter or transcripts. B. Comparison of H3K4me3 cfChIP-seq signal from a healthy donor (H012.1) with expected gene expression levels (Methods). Each dot is a gene. x-axis: normalized number of H3K4me3 reads in gene promoter. y-axis: expected expression in number of transcripts/million (TPM). C. Heatmap showing patterns of the relative H3K4me3 cfChIP-seq coverage on promoters of 14,875 RefSeq genes. The normalized coverage on the gene promoter (Methods) was log-transformed (log2(1+coverage)) and then adjusted to zero mean for each gene across the samples. The samples include cfChIP-seq samples from a compendium that includes healthy donors, acute myocardial infarction (AMI) patients, liver disease patients and CRC patients. D. Zoom in on the bottom cluster of (C). The right panel shows the H3K4me3 ChIP-seq from tissues and cell type from Roadmap epigenomics ^4^ and BLUEPRINT ^49^. Specific clusters of genes are marked by arrows. E. Genome browser view for megakaryocyte- and erythroblast specific genes. Shown is cfChIP-seq from two healthy samples (H012.1 and H013.1) and an AMI subject who exhibited enhanced erythropoiesis (M002.1). Also shown are two ChIP-seq profiles from the Roadmap Epigenetic reference atlas, and two samples from the BLUEPRINT project of cord-blood derived megakaryocytes and erythroblasts.

Next, we examined the relation between transcriptional levels and cfChIP-seq H3K4me3 signal. Comparison of H3K4me3 cfChIP-seq signal at promoters shows a good agreement with RNA levels in cells known to contribute to the cfDNA pool (*r*2 = 0.40; Figure 4B). This correlation is similar to a comparison between H3K4me3 ChIP-seq signal and RNA levels in matching tissues ^4^ (0.35 < *r*2 < 0.45, Figure S5A for an example), but not with that of an irrelevant tissue (Figures S5B and S5C).

Together, these results strongly suggest that H3K4me3 cfChIP-seq signal is informative of gene expression levels in tissues of origin.

### cfChIP-seq survey of diverse physiological and pathological conditions

Can cfChIP-seq profiles capture signals that reflect the underlying physiology? To better understand the variation of cfChIP-seq signal among subjects and in different physiological conditions, we performed H3K4me3 cfChIP-seq on 268 samples from a diverse cohort of subjects (clinical details summarized in Table S3). These include: 88 samples from ^61^ healthy donors (ages 23 - 66); 8 samples from four patients admitted to the emergency room with acute myocardial infarction (AMI); 38 samples from 33 patients with a range of liver-related pathologies; and 135 samples from 56 patients with metastatic CRC. There is expected variation in the cfDNA content among these patients due to changes in the contributing tissue of origin. For example, we expect to detect cfDNA from cardiomyocytes following AMI 27, cfDNA from colon tumors in CRC patients ^46,47^, and an increase in hepatocyte cfDNA in various liver pathologies ^30^.

To get a bird’s eye view of the differences in cfChIP-seq signal among samples, we performed a hierarchical clustering of 14,875 RefSeq genes promoters that have a noticeable signal in at least one sample across 268 samples (Methods). The clustering shows several trends (Figure 4C). A large group of 10,177 genes shows relatively small differences among samples. As expected, these genes tend to be highly expressed, housekeeping genes with CpG-island at their promoters (Figure S5D and S5E). The remaining 4,698 genes display a rich tapestry of patterns (Figure 4D). Strikingly, the variability among patients is much higher compared to healthy donors. We explore this variability in detail below.

Several clusters display high signal in healthy donors. Among these are clusters enriched for neutrophils (Cluster d) and for liver (Cluster b) that have observable signals in healthy donors, in agreement with previous studies ^16,31^. In contrast we see large clusters (Clusters a and c) enriched for GI tract and other solid tissues which show minimal signal in healthy donors.

### Platelet progenitor cfDNA in healthy donors

Our analysis identified a cluster with a positive signal in healthy donors (Cluster e, Figure 4D) that is enriched for megakaryocytes-specific genes such as GP6 and PF4 (25/144 genes in the cluster are in the REACTOME “Platelet activation, signaling and aggregation”, *p* < 2x10^-25^). However, there are no previous reports of megakaryocytes as a source of cell-free DNA. Conversely, previous analysis of cfDNA CpG methylation identified erythroblasts as major (20%-40%) contributors of cfDNA ^31,48^. These erythroblast-specific promoters are largely absent in healthy samples (Figure 4E). Two lines of evidence suggest that this is not due to the inability of cfChIP-seq to detect signals in these genes. First, H3K4me3 in ChIP-seq of erythroblasts and megakaryocytes derived from cord blood in the BLUEPRINT project ^49^ show the expected marking of the relevant genes in each cell type (Figure 4E). Second, there is a dramatic increase in cfChIP-seq signal of erythropoiesis related genes in one of our samples (e.g., GYPA, GYPB and ASHP, Figure 4E). This patient suffered from severe hypoxia at the time the blood was drawn and displayed signs of increased production of red blood cells (high RDW and low RBC and HGB; Table S3). Of note, this increase in erythropoiesis related genes is accompanied by a decrease in the signal for megakaryocyte-specific genes (Figure 4E).

Altogether our results suggest that platelet progenitors but not erythrocyte progenitors are major contributors to the cfDNA pool in healthy donors. The possible source of the discrepancy is lineage adjacency of erythrocytes and megakaryocytes who are both derived from a common hematopoietic progenitor ^50^, and thus may have similar CpG methylation patterns. Indeed, re-examining the DNA methylation analysis ^48^ using BLUEPRINT bisulfite sequencing data ^49^ we find that the region used is unmethylated also in megakaryocytes (Figure S5F), and thus it does not differentiate between these two cell types. This observation highlights the value of gene expression oriented information produced by cfChIP-seq in detecting events that are indistinguishable otherwise.

### cfChIP-seq detects cell of origin expression programs

To detect the compositions of cells/tissues that contribute to the cfDNA pool we defined cell-type/tissue-specific signatures from published ChIP-seq data4,49 (Figure 5A). We defined genomic locations (e.g., promoters) that have high signal only in the cell type in question (Methods, Table S4). Using this set of unique signatures we can test whether there is a contribution of the particular cell-type to the cf-nucleosome pool, since the only possible source of a signal in these loci is from that specific cell type. To detect the presence of a cell-type specific signature we compute the cumulative signal of the signature and contrast it against the null hypothesis of non-specific signal (Methods). In healthy donors, we observed a strong signal of neutrophil-, monocyte-, and megakaryocyte-specific signatures, and a lower but clear and significant signal of liver-specific signature, in agreement with published cfDNA methylation analysis ^31^ (Figure 5B). In contrast, no significant signatures from additional tissues such as heart and brain were observed (Figure 5B).

**Figure 5:**
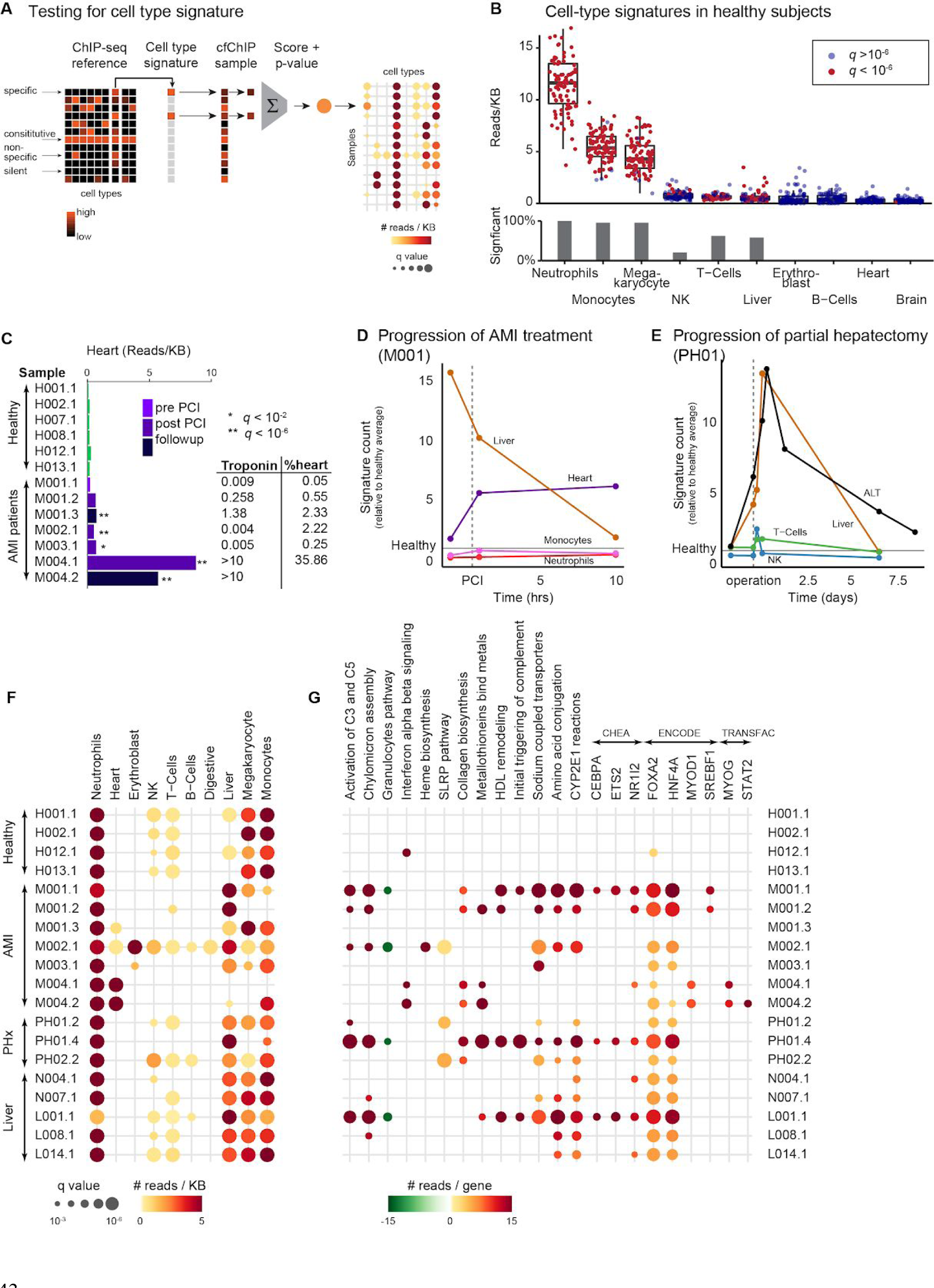
cfChIP-seq identifies cell-type specific and program specific expression patterns. A. Schematic outline of how we define and test cell-type specific signature. Using the existing compendium of ChIP-seq profiles, we define for each cell type a set of locations that are high only in the target cell type and low in all others. Given a cfChIP-seq profile, we sum the signal at all signature locations and test against the null hypothesis that this signal is due to non-specific background (Methods). B. Evaluation of average signal for signatures of different cell types in 88 healthy samples from ^61^ donors. Top: values (normalized reads/Kb) for each signature on all samples. Each dot is a sample, and boxplots summarize distribution of values in each group. Dots marked in red indicate values significantly different from background levels (Methods). Bottom: percent of samples with significant signal for each signature. We observe evidence for H3K4me3 signals in promoters specific to neutrophils, monocytes, megakaryocytes, NK cells, liver cells, and T-cells. We observe mild signals, but not significantly above background, from B-cells, and no signal from heart or brain. These proportions are consistent with independent estimates based on DNA CpG methylation marks ^31^. C. H3K4me3 cfChIP-seq signal in heart-specific locations in representative samples of healthy donors and acute myocardial infarction (AMI) patients (see Table S3 for details). **Inset**: measured troponin levels and percent cfDNA from cardiomyocytes as estimated using DNA CpG methylation markers ^27^ from the same blood draws. D. Changes in signature strength in an AMI (M001) patient before/after PCI. Signatures strength are normalized to the mean in healthy donors. Note the increase in heart and decrease in liver signatures post PCI. E. Changes in cfChIP-seq liver signature (brown line) and ALT levels (liver damage biomarker, black line) from blood samples of a patient that underwent partial hepatectomy (PH01). F. Heatmap showing significance of selected cell-type signatures in selected healthy donors and patients. See Table S5 for all samples and all signatures. Circle radius represents statistical significance (FDR corrected q-value) and the color represents read- density (normalized reads per kb) (Methods). G. Heatmap showing significance of selected gene sets from curated databases of transcriptional programs ^53^ and transcription factor targets ^81–83^ (Methods). See Table S6 for all samples and all signatures. The signal in each gene set is tested against the null hypothesis of levels similar to healthy donor baseline (Methods). Circle radius represents statistical significance (FDR corrected q-value) and the color represents the- average read number (normalized reads per genes) compared to healthy baseline (Methods).

As controlled test cases for cell-type detection, we considered pathologies where an increase in the signal of specific types of cells is expected. One such case is AMI, which involves the ongoing death of cardiomyocytes. A cardiomyocytes signal is not observed in healthy donors (Figure 5B), but clearly detected in samples from AMI patients undergoing percutaneous coronary intervention (PCI) (Figure 5C). Comparing our results to other measurements, we see good agreement between the strength of the cfChIP-seq heart signature, the levels of troponin measured in the blood, and the estimate of heart cfDNA by CpG methylation ^27^ (Figure 5C). When examining the changes in heart signature from admittance to the emergency room to post-PCI checkup (Figure 5D), we see an increase in heart signature immediately following the procedure, as previously reported by assaying cfDNA methylation ^27^.

Another test case involves recovery from partial hepatectomy where we expect to observe increased liver cell death. Indeed, we observed dramatic changes in the cfChIP-seq signal of liver signature following the operation which persisted for a few days and decayed to basal levels (Figure 5E). These changes are strikingly consistent with measurement of the liver marker ALT. A noticeable difference is the faster drop in the cfChIP-seq liver signal compared to ALT, likely reflecting the shorter half-life of cfDNA (<1 hour) relative to ALT (∼47 hours) in the circulation ^51^.

An important advantage of cfChIP-seq is that it is not limited to a set of preselected markers and hence it can provide an unbiased view of the contributions of different cell types to the cfDNA pool. To implement such an unbiased approach, we evaluated the panel of cell-type specific signatures across all of our cfChIP-seq samples (Figure 5F and Table S5). This analysis shows that in all samples we can detect signatures of leukocytes (e.g., monocytes and neutrophils), and remote organs (e.g., liver and bone marrow megakaryocytes). Of note, the observed decrease in the relative level of leukocyte signatures in samples that show increased cfDNA load, is consistent with a smaller proportion of cfDNA from these cells. For example, AMI patient M004.1 had cfDNA concentration of 21ng/ml and 35% his cfDNA originated from heart based on CpG methylation analysis.

This unbiased approach reveals a more complex picture in AMI patients. In addition to the heart signature discussed above, in some AMI patients we observe a significant increase in cfChIP-seq liver signature both before and shortly after PCI (Figure 5D). This signature includes a clear signal at liver-specific genes, such as Albumin and complement genes (Figure S6B). This increase is presumably due to liver injury in AMI patients secondary to low organ perfusion and liver hypoxia ^52^. To confirm the unexpected liver cfChIP-seq signal in AMI patients, we analysed the cfDNA methylation status for liver-specific DNA methylation regions indicative of liver cell death ^30^, and find excellent agreement between liver cfChIP-seq signature levels and liver cfDNA estimates (R2=0.96, Figure S6C).

Together, these results demonstrate that cfChIP-seq signal reflects differences in the tissue of origin composition. In particular, where ongoing pathological processes take place, cfChIP-seq signal corresponds to the affected tissue, such as heart and liver.

### cfChIP-seq signal reflects patient-specific transcriptional programs activity

Since cfChIP-seq signal correlates with the gene expression programs in the cells of origin, we proceeded to inquire whether cfChIP-seq can reveal specific transcriptional programs within the tissue of origin. To test this hypothesis, we evaluated the H3K4me3 cfChIP-seq signal in gene sets representing different cellular processes, protein complexes, transcriptional responses based on gene expression studies, and targets of transcription factors based on ChIP studies ^53–56^ (Figure 5G, Methods). Many of these sets include genes that are not unique to a certain cell type and are specifically marked by H3K4me3 in cells that contribute to the cf-nucleosomes pool in healthy donors. We thus devised a test for changes in the signal of a gene set (sum over all the genes within the set) compared to the mean and variance of a reference healthy cohort of ^26^ samples (Methods). This analysis uncovered multiple gene sets which signal differs from the expected signal --- that is, the amount of H3K4me3 cfChIP-seq signal for the gene set in a subject is significantly different from the observed signal in the reference healthy cohort (Table S6).

cfChIP-seq identifies patient specific transcriptional programs. For example, in M002.1 we observe a strong increase in the signal of Heme Biosynthesis (*q* < 10-9) and a strong decrease in Granulocytes Pathway (*q* < 10-9), consistent with the results discussed above (Figure 5D). Another example is the increased interferon signature in M004, who suffered a severe heart damage as reflected by the levels of troponin and cfChIP-seq heart markers (Figure 4C). Induction of interferon response was recently shown to promote a fatal response to AMI ^57^. The induction of interferon-mediated immune response is accompanied by increased cfChIP-seq signal in targets of STAT2 and other immune-related transcription factors. In addition, consistent with the massive amount of cardiomyocyte cfDNA in M004, we observed a significant increase in targets of MYOD1 and MYOG1, two factors involved in cardiomyocyte development.

### Detection of pathology-specific liver signals

The dynamic nature of active histone marks suggested the hypothesis that cfChIP-seq may inform on intra tissue pathology-related alterations in gene expression. To test this hypothesis, we decided to focus on hepatocytes since 1. Hepatocytes play a central role in multiple facets of physiology - metabolism, metabolite storage, protein synthesis and degradation, blood homeostasis, bile production, and drug clearance and 2. Many of the gene programs enriched in our samples are related to liver function (e.g., HNF4A and FOXA2 targets, triggering of complement, and cytochrome P450 complex; Figure 5G). We assumed that differences in disease etiology and presentation could reflect in different cfChIP-seq signals despite common tissue of origin. We first assembled a cohort of subjects with verified liver-related diagnosis and/or subjects showing increased liver contribution. This cohort included subjects at different stages of Nonalcoholic fatty liver disease/Nonalcoholic steatohepatitis (NAFLD/NASH) (n=15), autoimmune hepatitis (AIH) (n=3), post liver transplant (n=5), infection associated with liver injury (n=1), AMI-associated liver injury (n=1) and partial hepatectomy patients (n=2) as discussed above (Table S3).

We estimated the percentage of liver-derived chromatin in each sample using the Roadmap Epigenomics liver H3K4me3 ChIP-seq sample as an instance of pure liver tissue (Figure 6A, Methods). The resulting estimates range from ∼2% in healthy samples to 44% in liver patients. These estimates are consistent with the previously reported 2-4% liver contribution in most healthy donors ^31^ and with a CpG methylation based estimate of liver cfDNA quantity (*r2* = 0.87, Figure S6D).

**Figure 6:**
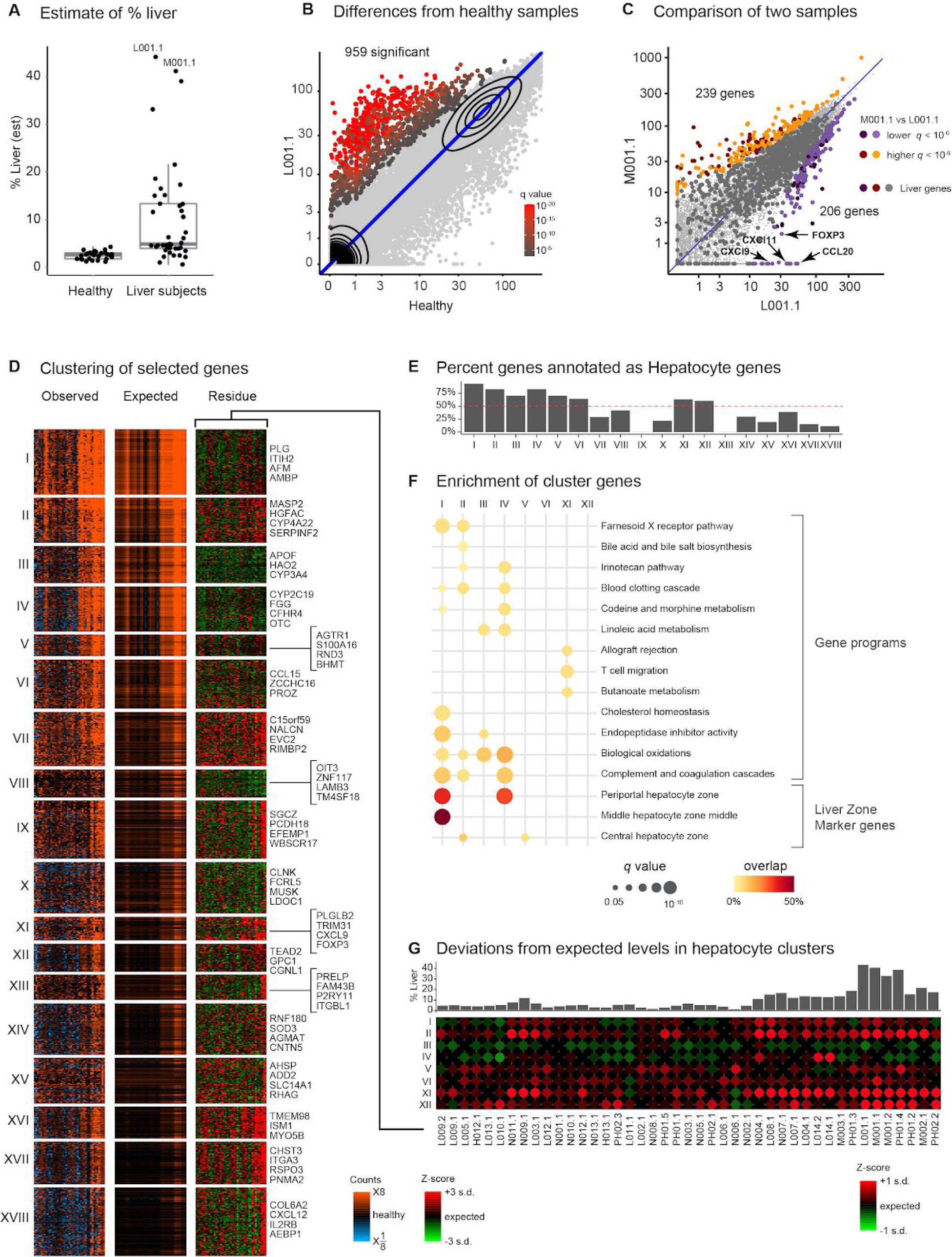
cfChIP-seq detects changes in liver-specific transcriptional programs. A. Estimate of %liver contribution to healthy reference cohort and a cohort of subjects with various liver-pathologies (Table S3). B. Evaluation of differentially marked genes in a sample of an acute autoimmune induced hepatitis subject (L001). For each gene we compare the mean normalized coverage of a healthy reference cohort (x-axis) against the normalized coverage in the sample (y-axis). Color indicates whether the observed number of reads is significantly higher than we would expect based on the distribution of values in healthy samples (Methods). C. Differentially marked genes between two samples with similarly high liver contribution L001.1 (acute AIH) and M001.1 (AMI induced liver damage). For each gene we compare coverage in the two samples (L001.1, x-axis; M001.1 y-axis). Significance test whether the two values are sampled from the same distribution (Methods). Dark circles - genes that are significantly different in liver ChIP-Seq (Roadmap Epigenomics) compared to healthy reference. D. Clustering of 1,320 genes that are significantly higher in one of the samples in the liver cohort compared to healthy samples baseline. Left panel: values compared to healthy baseline. Middle panel: expected level assuming healthy liver signal and %liver contribution. Right panel: Z-score of observed value from expected (mean and variance) value. For each cluster we show 3-4 representative genes (right). Sample order in each heatmap is identical and matches the order in (G). E. Percent of genes in each cluster of (D) that are annotated as hepatocyte genes ^60^. Clusters above the 50% threshold (red dashed line) are considered as hepatocyte origins. F. Enrichment analysis of hepatocyte clusters (Clusters I-VI, XI, and XII). Hypergeometric test for significant overlap with gene programs from curated databases ^84^ and marker genes of hepatocyte zones ^61^. Circle radius: q-values of hypergeometric enrichment test, circle color - fraction of overlap. G. Top: Percent of liver contribution in each sample in the order shown in (D). Bottom: Deviations from expected values for each sample in each of the hepatocyte clusters (average Z-score for each sample on cluster genes).

For example, sample L001.1 is taken from a young child suffering from AIH. We estimated that 44% of its cfDNA is liver-derived. Comparing the signal of L001.1 to healthy reference we find 959 genes with significantly increased cfChIP-seq signal (Figure 6B). These genes are highly enriched for liver functions and hepatocyte genes (*q* < 10-250).

To understand whether this increase in liver genes signal is universal to all liver pathologies, we compared L001.1 with M001.1, a sample from an AMI patient that has similar estimated levels of liver contribution (41% liver). As expected, many liver-specific genes are similarly increased in both samples (Figure 6C, dark gray circles). We did not, however, observe pronounced differences between the two samples (Figure 6C) with hundreds of genes significantly higher in only one of these samples. In L001.1 we observe enrichment for genes involved in interferon gamma signaling (*q* < 2x10^-8^), immune system (*q* < 1.2x10^-7^), MHC class II protein complex (*q* < 1.4x10^-6^), and allograft rejection (*q* < 1.3x10^-5^), consistent with the autoinflammatory state of this patient. We also detect stronger signal in genes associated with AIH such as the transcription factor FOXP3, and the interferon gamma induced chemokines CXCL9, CXCLl1, and CCL20 ^58,59^ (Figure 6C). Importantly, several of these genes (dark colors) are liver specific, demonstrating a potential of cfChIP-seq in detecting intraorgan transcriptional changes. The liver-specific genes that show specific increase in L001.1 are enriched for genes involved in complement and coagulation (*q* < 1.5x10^-4^), such as CFH and C4BPA, consistent with the immune phenotype in this patient. M001.1 shows relative increase in genes enriched for neutrophil-mediated immunity (*q* < 5.8x10^-8^). This could also be due to a decrease in this signal in L001.1. Focusing on liver-specific genes we see enrichment for metabolism-related pathways such as metabolism of xenobiotics by P450 (*q* < 1.1x10^-5^), and bile secretion (*q* < 1.5x10^-4^).

To get a more systematic view of the differences between samples that are due to changes in liver-specific expression programs rather than the amount of liver contribution, we focused on 1,320 genes whose expression is significantly higher than expected in one of the liver cohort samples, compared to the healthy cohort, (Figure 6D, left panel). We calculated the expected signal per gene based on the estimated liver contribution of the specific sample and the ChIP-seq signal of the gene in healthy liver tissue (Figure 6D, middle panel, Methods). We next used the expected and observed values to calculate a Z-score --- the extent of deviation of the observed signal from the expected value, accounting for both sampling noise and the variability observed between healthy donors (Figure 6D right panel). We then used this score to cluster the matrix (Methods, Figure 6D) .

This analysis identified gene clusters for which the expected signal explains most of the variation between samples (e.g Clusters I, III, and IV), suggesting that most of the signal in these clusters is due to contribution from liver cells. In other clusters, such as Cluster XV the signal is not explained by the amount of liver contribution and indeed, many of the genes in the cluster are expressed specifically in erythrocyte progenitors (e.g., ASHP and HBD; 37/78 genes, *q* < 10-12, Figure 6D). In some clusters, such as Clusters II, V, and XI, the amount of liver contribution explains some of the observed differences, but not all of them. These genes can be either differentially expressed in the liver in some of the subjects, or originate from a mixture of several different tissues (e.g., liver and heart). To better understand the contribution of liver-specific transcriptional programs, we focused on clusters where at least 50% of the genes are annotated as hepatocyte genes ^60^ (Figure 6E, Clusters I-VI, XI, XII). Such analysis may identify additional genes that are only expressed in abnormal hepatocytes, due to coexpression with healthy hepatocytes genes within the cluster..

Next we performed enrichment analysis of the gene sets in each cluster (Figure 6F). As expected we see strong enrichments for many liver related terms (Table S7). Interestingly, some clusters show strong enrichments only to specific terms. For example, the genes of Cluster I are enriched for genes involved in the process of cholesterol homeostasis (9/111 genes, *q* < 4x10^-8^) and the genes in Clusters I and IV are enriched with genes of the complement and coagulation cascade (14/111 genes, *q* < 3x10^-15^, and 11/77 genes, *q* < 2x10^-12^, respectively). c

To further characterize the clusters, we examined a recent single-cell RNA-seq atlas of human liver cells ^61^. This atlas includes lists of marker genes for hepatocytes at different liver zones which represent division of labor among hepatocytes in the axis from the portal vein, (input to the liver from the gastrointestinal tract) to the central vein (output from the liver) ^62^. Testing our gene clusters against these marker genes we see that Clusters I and IV are enriched for marker genes of periportal hepatocyte zones, Cluster I is also enriched for genes of middle hepatocyte zones, and Clusters II and V are enriched for marker genes of central hepatocyte zone (Figure 6F). These could indicate either increased cell death in the relevant zone, or global changes in liver metabolism toward the relevant metabolic regime.

This analysis demonstrates that unbiased clustering of the data captures meaningful functional modules of hepatocyte biology. Examining the deviations in the signal of clusters between samples allows us to identify samples-specific changes in hepatocyte-specific transcriptional programs (Figure 6G). For example, we see high levels of Cluster I genes in patients with immune response (L001 - acute AIH; L004 - chronic AIH; L008 - liver transplant; L014 - localized inflammation; and some of the NASH patients N004). In contrast, we see high levels of Cluster IV genes in a subset of these patients (N004 and L014). Thus, although these clusters are both enriched for the periportal zone markers (Figure 6F) they capture transcriptional programs that are differential among subjects in the liver cohort.

Together, these results demonstrate the ability of cfChIP-seq to detect cell states within a remote tissue (liver) and within a specific cell type (hepatocytes).

### Analysis of colorectal-cancer by cfChIP-seq

To test if cfChIP-seq can provide information on tumors, we analyzed a collection of samples from an ongoing longitudinal study following metastatic CRC patients before and during treatment, including patients with undetectable or minimal disease at the time of sampling. Overall we analyzed 193 samples from 67 patients, of which ∼70% (135/193 samples from 56 patients) passed our QC criteria. We see this rate as encouraging since blood draws, samples collection and storage of these samples was not designed for cfChIP-seq analysis.

As observed above (Figure 4C) samples from within the CRC cohort showed much higher cfChIP-seq signal variability than among healthy donors. Indeed, examining the pairwise correlations within each cohort of samples highlights the differences between healthy donors and cancer patients (Figure 7A): Closely collected samples show higher similarity than samples collected far apart suggesting that to a large extent the variability among cancer samples is due to differences in the underlying patient molecular state ^63^. To detect CRC-specific signals, we generated two signatures, a “digestive signature” based on the Roadmap digestive tissue ChIP-seq, and a “COAD signature” based on analysis of the TCGA gene expression data of colorectal adenocarcinoma (COAD) samples (see methods). Using these signatures, that are derived completely from external data, we correctly classified CRC samples with AUC of 0.83, and 0.84 for digestive and COAD, respectively (Figure S7A). Examining the value of these two signatures in all samples, we observed a large variation among the cancer samples (Figure 7B) and low values in healthy donors.

**Figure 7:**
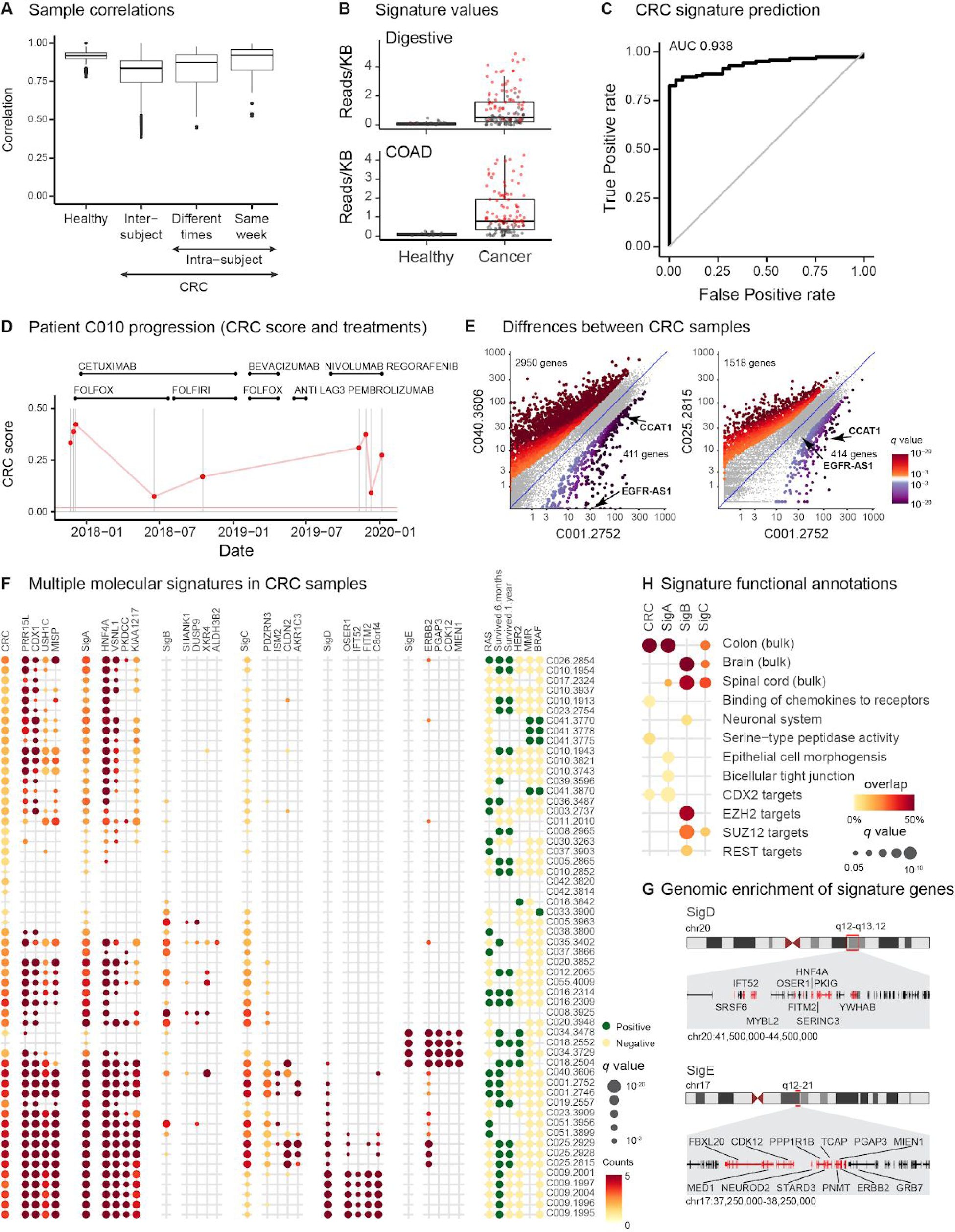
cfChIP-seq identifies molecular heterogeneity in colorectal carcinoma patients. A. CRC samples are much more variable than samples of healthy donors. Box plots show correlation (pearson correlation, y-axis) of pairwise comparisons: between healthy donor samples; between CRC samples from different patients; between CRC samples of the same patient taken more than a week apart; between CRC samples of the same patient less than a week apart. B. Distribution of signature strength of healthy and CRC samples. Top: signature of digestive tissue (as in Figure 5F). Bottom: COAD gene signature. Box plots show distribution of signal (Reads/KB, y-axis) in each group. Each sample is a dot, red = significantly above background (Digestive) or healthy baseline (COAD). C. Classification accuracy of CRC patients vs healthy donors. Shown are fraction false positive (x-axis) vs fraction true positives (y-axis) for different thresholds of CRC signature. Diagonal line: expected curve for random classifications. D. Progression of cancer signature during treatment of a single patient. Top: treatment history of the patient as a function of time (x-axis) Bottom: CRC signature strength (y-axis) for different samples. E. Differences between CRC samples with high CRC signature strength. For each gene we compare coverage in the two samples (x-axis and y-axis). Significance test whether the two values are sampled from the same distribution (Methods). F. Evaluation of cancer signatures on samples with high tumor percent. Shown is the CRC signature (based on TCGA genes) and five signatures found by our analysis. For each signature we show the level of the signature and of four representative genes. Circle color represents the increase in counts/gene above healthy reference samples, and circle radius represents significance of this increase. Rightmost panel displays major clinical parameters of the sample: RAS, BRAF mutations, HER2 amplification, MMR deficiency, and survival above 6 month and 1 year after the sample was taken. G. Functional enrichment of signatures. Shown are representative enrichment from an unbiased evaluation of signature genes against large annotations database (EnrichR ^84^. See Table S9 for the complete set of enrichments. H. Genome clusters containing SigD and SigE signatures. Marked in red are genes from each signature in the specific genomic loci. (Names of noncoding transcripts were omitted).

These results suggest that there is a large variability in the amount of tumor-derived cf-nucleosomes among the samples. To estimate the tumor-related contributions we selected a subset of COAD-genes that are not observed at all in a reference cohort of healthy donors and used them as a “CRC” signature (189 genes). Assuming that the samples with the highest cancer signal have close to 100% tumor contribution, we calibrated these scores to the range of 0-1 representing a rough proxy of tumor load. Not surprisingly, the CRC signature strength is highly correlated with the digestive and COAD signatures discussed above, yet it has better predictive value (AUC = 0.94, Figure 7C).

We observed large differences in the CRC signature magnitude between patients and during treatment of the same patient, consistent with the course of therapy (Figure 7D) ^63^. In addition to changes in the CRC signature magnitude, we detect differences that appear to result from disease progression (Figure S7B). Some of the differences are due to increased liver signal in C010.1943 vs. C010.3743 (ARCHS4 tissue *q* < 3x10^-16^, which might reflect chemotherapy-induced liver damage ^64^. Other changes may reflect intratumor variation, or immune-related signaling such as the enrichment for interferon gamma genes in C010.3743 vs. C010.1943 (REACTOME *q* < 3.6x10^-6^, Figure S7B).

### cfChIP-seq detects molecular variability among colorectal-cancer patients

A hallmark of cancer cells is genetic alterations that lead to dysregulated gene expression programs ^65^. Identification of such cancer-specific transcriptional programs can assist treatment choice ^66^. We asked whether cfChIP-seq can reveal transcriptional programs associated with the disease.

A comparison of samples from different patients with similar CRC signature levels revealed striking differences in hundreds of genes (Figure 7E). These differences can be due to contribution of additional tissues (e.g enrichment for liver genes in C001.2752 vs. C040.3606, ARCHS4 tissue *q* < 10-9), while others may reflect intertumor transcriptional differences for example enrichments for Wnt/calcium/cyclic GMP pathway in C040.3606 vs. C001.2752 (BioPlanet q<0.00025) and for Cell adhesion molecules (CAMs) in C025.2815 vs. C001.2752 (BioPlanet q<10-4). Additional examples include, EGFR-AS1, and the CRC marker CCAT1 ^67^ (Figures 7E and S7C). EGFR-AS1 regulates the splicing of EGFR and may affect anti EGFR treatment ^39^. Interestingly, when examining all samples, we identify variation in genes associated with immune activity such as the checkpoint receptors CD160, TIGIT, and PDL1 (CD274) (Figure S7D), suggesting that we may detect tumor-related immune signals.

To identify major cfChIP-seq signature subtypes, we tested the gene set compendium (discussed above) against samples with relatively high cancer load (56 samples from ^29^ patients, where CRC Signature > 0.15). We found 680 (out of 7,538) gene sets that had informative signals in these samples (Table S8, Methods). We used these to initialize an iterative process to identify signatures that distinguish between samples subgroups (Methods) resulting with five gene signatures that capture the main behaviors in the original set of programs (Figures 7F). Signatures A-C capture cancer gene expression programs and signatures D-E capture duplications events.

The scores of the largest signature (SigA) are highly correlated with the CRC scores, although there is only a partial overlap between the two (Figure S7F). The genes in this signature are enriched with genes associated with Colon (ARCHS4 tissue *q* < 10-64), targets of CDX2 a transcription factor active in CRC (TRRUST, *q* < 10-9) (Figure 7H and Table S9). The second signature (SigB) differentiates a small subset of the high CRC samples. Interestingly, this signature is enriched for genes in neuronal associated terms (Brain, ARCHS4 tissue *q* < 10-39). The genes in this cluster are also enriched for Polycomb Repressive Complex (PRC) and REST targets (ENCODE and ChEA, SUZ12 *q* < 10-22, EZH2 *q* < 10-22, REST *q* < 3.7 x 10-8). REST represses neuronal genes in colon epithelium, and is often deleted in CRC tumors ^68^. This could indicate derepression due to loss of polycomb/REST activity leading to misregulation of neuronal genes in the tumor. Alternatively, it may indicate involvement of neuronal phenotype in these tissues ^69^. The third signature (Sig C) selects a larger subset of samples, which includes most of the samples selected by SigB although there is little overlap of genes between the two signatures (Figure S7F).

To examine the clinical significance of the expression signatures A-C we compared them to the Consensus Molecular Subtypes (CMS) classification of CRC tumors ^70^. We examined the behavior of these signatures in 198 labeled CRC tumor gene expression profiles in the TCGA database ^41^ (Figure S7G). This analysis shows that SigA genes tend to have lower expression in CMS1 tumors, while SigB genes tend to have higher expression in CMS4. CMS1 tumors are characterized by genome instability, increased immune infiltration and immune response activation. CMS4 tumors are characterized by upregulation of epithelial to mesenchymal transition (EMT) and cancer stem cell like phenotype and have been shown to have low EZH2 expression ^71^. This characterization of CMS4 is consistent with the REST and PRC de-repression observed in SigB (Figure 7H).

Ten out nineteen genes in Signature D and 13 of the 17 genes in signature E are clustered around regions of known genomic duplications at chr20q13.12, and chr17q12-q21, respectively (Figure 7G) ^72,73^. The chr20q13.12 amplification has been previously reported in CRC and includes HNF4A, a transcription factor with increased activity in CRC ^72^. The chr17q12-q21 includes the gene ERBB2, and known as the HER2 amplicon that appears in multiple types of cancer and has prevalence of 4% in CRC ^72^. Consistently, SigE is high in samples with identified HER2 amplifications (Figure 7F), suggesting that cfChIP-seq detects this massive genomic amplification event. Unlike genomic copy number, which can be detected using background reads (Supplemental Note), the H3K4me3 cfChIP-seq signal further increases the confidence that these copy number variations involve active transcription in the amplified regions. Detection of HER2 amplification in colon cancer has significant practical implications since it is a predictive marker for prolonged survival of patients treated with HER2 inhibitors^74^. This intriguing potential requires future validation on a larger cohort.

Altogether these results show that a single cfChIP-seq blood test have the potential to detect the variability in CRC patients related to the load of the tumor (CRC score), the contribution of additional tissues (e.g., liver damage, immune cells), and gene expression inter-tumor heterogeneity. The latter feature of CRC is uniquely revealed by cfChIP-seq.

## Discussion

Here we introduce cfChIP-seq to infer the transcriptional programs of dying cells by genome-wide mapping of plasma cf-nucleosomes carrying specific histone marks. We demonstrate the feasibility to perform ChIP-seq on plasma cell-free nucleosomes with four histone marks associated with active transcription (H3K4me1, H3K4me2, H3K4me3, and H3K36me3) for probing active or paused enhancers and promoters, and gene body-associated transcriptional elongation. We further performed in depth promoter centric analysis on a large cohort of ^61^ healthy donors, and 89 patients, including 135 samples from metastatic colorectal cancer patients. This analysis shows that cfChIP-seq can detect signals beyond the resolution of cells of origin. We show differences in hepatocytes-specific transcriptional programs between subjects with different etiology of increased liver cfDNA (Figure 6). Our analysis shows that even at this early stage, cfChIP-seq is highly sensitive in detecting signatures of interest, including cancer-specific signatures (Figures 3 and 7). Since the sensitivity of cfChIP-seq is increased with the signature size, we anticipate that the potential sensitivity can be significantly increased by using multiple antibodies that target different genomic landscapes. Importantly, the assay leaves most of the original sample intact, allowing reuse for multiple assays which is important in many clinical situations where blood volume is a limiting factor. A unique feature of cfChIP-seq is that the immunoprecipitation step generates a biologically relevant reduced representation of the genome. This allows us to perform genome wide unbiased analysis without the need for preselecting markers and with low sequencing depth. This can be important for detection of changes in tumor makeup following treatment, and may allow simple patient-specific analysis for the detection of minimal residual disease, at low cost.

Most current cfDNA-based methods rely on detecting genomic alterations in cfDNA to quantify the contribution of cfDNA from cells with altered genomic sequence, such as a fetus, a transplant, or mutated genes in tumors ^15–17,23^. These methods are blind to events that involve turnover and death of somatic cells. More recent approaches leverage epigenetic information in cell free DNA. Extremely deep sequencing of total cfDNA to identify nucleosomes and transcription factors positions ^19,75^ and occupancy ^18^ reflect tissue of origin and gene expression. However, they rely on detecting changes in coverage over target regions, with a signal of each tissue/cell type imposed on the background of all other tissues/cell types (e.g., detection of an event causing nucleosome depletion in 1% of the cells requires distinguishing the difference between 99% occupancy and 100% occupancy). Even with extremely deep sequencing coverage (100s of million reads per sample), there is a prohibitive harsh detection limit for events in rare subsets of cells ^19^. Recently, machine learning methods were used to detect several types of cancers by analysis of fragment lengths with shallower cfDNA sequencing 32, achieved by focusing solely on the classification task without detection of specific cell-types contributing to cfDNA. An alternative modality is assaying cfDNA CpG methylation along the sequence ^20–22,25,27–31^. DNA methylation serves as a stable epigenetic memory and is largely unchanged upon dynamic cellular responses. As such, it is highly informative regarding cell lineage, but much less about transient changes in expression.

Many cellular processes, including cancerous transformation involve large changes in transcriptional programs, which leave unique imprints on the histone modification landscape. Intensive research during the last two decades established an intimate connection between specific histone marks and transcription. Specifically, the levels of H3K4me3 correlate with gene expression (Figures 2C and 4A), which allow us to infer transcriptional activity in cells of origin. Therefore, assaying chromatin marks in cf-nucleosomes provides rich and complex information beyond current methodologies. Furthermore, the advent of single-cell technologies is constantly uncovering new cell types, cell states, and their specific molecular biomarkers ^76^. cfChIP-seq opens an opportunity to tap these insights in non-invasive liquid biopsy.

We exploit the wealth of knowledge about gene expression for interpreting cfChIP-seq results. For example, observation of cfChIP-seq signal from genes encoding platelet-specific proteins (e.g GP6, GP9), but not erythrocyte-specific proteins (e.g., HBB) in healthy donors led us to identify megakaryocytes but not erythroblasts as major cfDNA contributors in healthy donors. Another example is our analysis of liver pathologies. By using existing annotations (all based on gene expression studies) we identified the genes that represent hepatocyte contribution to the signal. We then used marker genes identified in a recent liver single cell RNA-seq atlas ^61^ to detect different contributions from different liver zonation expression programs in each of the subjects. Finally, in our analysis of the CRC cohort we used a large collection of gene sets, mostly from gene expression studies ^53^ as the basis for identifying signatures that classify molecular phenotypes of the samples.

These examples demonstrate the potential of using a single histone mark focused at gene promoters. There are potential advantages to combine multiple chromatin marks. Using H3K36me3 cfChIP-seq, which marks active elongation we can better distinguish between a poised state and actual transcription. Parallel analysis of enhancer chromatin marks such as H3K4me1/2 can provide more precise understanding of the regulatory program that activated the genes. It is often the case that the same gene is regulated by multiple enhancers that are responsible for its activation in a specific cell type or specific transcriptional response. Thus, observation of enhancer activity provides wider context to the activation of its target gene(s). The main challenge in harnessing this information is our partial knowledge of enhancer-gene interactions in multiple tissues. This is the subject of much research and progress is made at a rapid pace ^77^.

More broadly, cfChIP-seq facilitates the systematic study of multiple chromatin marks in cf-nucleosomes. Beyond transcription, chromatin state is also intimately related to other chromatin-templated processes such as cell cycle progression and DNA damage and repair. The potential for observing such processes with a non-invasive assay can revolutionize our understanding of basic questions in human physiology and pathology. Our results establish the method and demonstrate its ability to probe the active and poised genes in cells of origins. To fully harness the potential of this assay we need a deeper understanding of the processes of cell death in health and specific pathologies. Improving our ability to bridge the gap between epigenomics and transcriptional states will allow us to better exploit the massive transcriptional profiles collected in different cell types, developmental stages, and pathological states for interpretation of cfChIP-seq profiles. Finally, by their nature, cell-free assays examine the superimposed contributions of multiple cell populations. Thus, unmixing, or deconvolving signals is a central challenge for improved interpretation ^78^.

Assaying modified cf-nucleosomes, either used alone or in combination with existing biomarkers, has multiple potential medical applications. We envision identifying not only the cells that are dying, but also the molecular basis for the pathological or physiological states. This may lead to a better understanding of complex diseases in which several cell types interact to elicit a pathology. An important example is the contribution of specific subpopulations of cells (e.g exhausted T-cells) residing in the tumor to the oncogenic process and success of treatment. This sub-population is defined by their transcriptional state, not their DNA sequence nor their cell lineage identity, and thus cannot be identified by current cfDNA methodology. Moreover most of these cells are restricted to the tumor, and thus cannot be easily detected by observation of the circulating immune cells. However, they turnover within the tumor, and thus can be detected in the cf-nucleosome pool (Figure S7D). Detection and monitoring of immune response within a remote organ has exciting implications. For example, more accurate monitoring of treatment response, leading to more precise administration of treatment in the short term, but also longer term insights at the transcriptional level of drug action mechanisms and side effects.

Altogether, cfChIP-seq is a highly informative and minimally invasive assay which opens up a wide range of opportunities for studying basic questions in human physiology that have been inaccessible until now.

## Acknowledgements

We thank N. Kaminski, J. Moss, E. Pikarsky, O.J. Rando, N. Rajewsky, A. Regev, and members of the Friedman lab for discussions and comments on this manuscript. We thank L. Friedman for help with illustrations and graphics. This work was supported by: European Research Council’s AdG Grants “ChromatinSys” (NF) and “RxmiRcanceR” (EG); Israel Science Foundation’s I-CORE program grant 1796/12 (TK and NF) and grants 2612/18 (NF); NIH Grants RM1HG006193 (NF); and Israel Innovation Authority’s Kamin grant 63381 (NF). EG is also supported by a number of ISF grants, Israeli MOS grant and an NIH grant. A patent application for cfChIP-seq has been submitted by the Hebrew University of Jerusalem.

## Materials and Methods

### Patients

All clinical studies were approved by the relevant local ethics committees. The study was approved by the Ethics Committees of the Hebrew University - Hadassah Medical Center of Jerusalem. Informed consent was obtained from all subjects or their legal guardians before blood sampling.

### Sample collection

Blood samples were collected in VACUETTE® K3 EDTA tubes, transferred immediately to ice and 1X protease inhibitor cocktail (Roche) and 10mM EDTA were added. The blood was centrifuged (10 minutes, 1500 × g, 4°C), the supernatant was transferred to fresh 14ml tubes, centrifuged again (10 minutes, 3000 × g, 4°C), and the supernatant was used as plasma for ChIP experiments. The plasma was used fresh or flash frozen and stored at -80°C for long storage.

### cfChIP-seq

#### Bead preparation

50μg of antibody were conjugated to 5mg of epoxy M270 Dynabeads (Invitrogen) according to manufacturer instructions. The antibody-beads complexes were kept at 4°C in PBS, 0.02% azide solution.

| AB | Company | Catalog Number |
| --- | --- | --- |
| IgG | Cell signalling | 2729S |
| H3K4Me1 | Diagenode | C15410194 |
| H3K4Me2 | Diagenode | C15410035 |
| H3K4Me3 | Diagenode | C15410003 |
| H3K36Me3 | Diagenode | C15410192 |

#### Immunoprecipitation, NGS library preparation, and sequencing

0.2mg of conjugated beads (∼2μg of antibody) were used per cfChIP-seq sample. The antibody-beads complexes were added directly into the plasma (1-2 ml of plasma) and allowed to bind to cf-nucleosomes by rotating overnight at 4°C. The beads were magnetized and washed ^8^ times with blood wash buffer (BWB: 50mM Tris-HCl, 150mM NaCl, 1% Triton X-100, 0.1% Sodium DeoxyCholate, 2mM EDTA, 1X protease inhibitors cocktail), and three times with 10mM Tris pH 7.4. All washes were done with 150ul buffer on ice by shifting the beads from side to side on a magnet. Do not use vacuum to remove supernatant during washes in buffers that do not contain detergents.

On-beads chromatin barcoding and library amplification was done as previously described ^33,34^ except for the DNA elution and cleanup step where the beads were incubated for ^1^ hour at 55°C in 50μl of chromatin elution buffer (10mM Tris pH 8.0, 5mM EDTA, 300mM NaCl, 0.6% SDS) supplemented with 50 units of proteinase K (Epicenter), and the DNA was purified by 0.9 X SPRI cleanup (Ampure xp, agencourt). The purified DNA is eluted in 25 μl EB (10mM tris pH 8.0) and 23 μl of the eluted DNA were used for PCR amplification with Kapa hotstart polymerase (16 cycles). The amplified DNA was purified by 0.8 X SPRI cleanup and eluted in 12 μl EB. The eluted DNA concentration was measured by Qubit and the fragments size was analyzed by tapestation visualization. Note: If adapter dimers are substantially visible by tapestation post library amplification, we recommend pooling samples and performing additional X 0.8

SPRI DNA cleanup, or separating the pooled samples on a 4% agarose gel (E-Gel® EX Agarose Gels, 4%, Invitrogen), and gel purification of fragments larger than adapter dimers (>150bp). DNA libraries were paired end sequenced by Illumina NextSeq 500.

#### Sequence analysis

Reads were aligned to the human genome (hg19) using bowtie2 (2.3.4.3) with ‘no-mixed’ and ‘no-discordant’ flags. We discarded fragments with low alignment scores (-q 1) and duplicate fragments.

See Table S1 for read number, alignment statistics, and numbers of unique fragments for each sample.

#### Roadmap Epigenome atlas

We downloaded aligned read data from the Roadmap Epigenome Consortium database (Table S10). For our analysis we discarded pre-natal, ESC, and cell-line samples, resulting with 64 tissues and cell types (Table S11). The aligned read files were then processed with the same pipeline as cfChIP-seq samples. That is, all steps from numbers of reads mapped to each genomic window, background estimation, normalization, etc.

#### Tumor-type Gene Signatures

We downloaded RNA-seq data from the TCGA and GTEx projects as analyzed by the Xena project ^79^ (Table S10). We defined the set of genes that are over-expressed in a tumor type to satisfy three requirements: 1) Significantly higher expression in tumor samples compared to the corresponding tissue samples (t-test, *q* < 0.001 after FDR correction); 2) Significantly higher expression compared to all healthy samples (t-test, *q* < 0.001 after FDR correction); and 3) Median expression in the tumor is higher than the median expression in each of the healthy samples.

#### Expected healthy expression level

To best emulate expression profiles of healthy individuals in the analysis of Figure 2C, we performed *in silico* mix of the four cells types that contribute the most to cfDNA 31:31: neutrophils, 32%; monocytes 32%; megakaryocytes 20%; and NK cells 5%. The gene expression for these cell types was downloaded from BLUEPRINT consortium website (Table S10).

#### TSS location catalogue

We downloaded the Roadmap Epigenome Consortium ChromHMM annotation of all consolidated tissues (Table S10). Using these annotations we constructed a catalogue of potential functional sites (enhancers, TSSs, and genes). We extended the catalogue to include 3kb regions centered on TSS of annotated transcripts in the UCSC gene database and ENSEMBL transcript database (Table S10). We used the combined catalogue to define regions along the genome. We used a different version of the catalogue for analysis of each antibody, to match the mark. For H3K4me3 analysis we used only TSSs, for H3K36me3 analysis we used only gene bodies, and for H3K4me2 we had annotations of TSSs and enhancers. In each version of the catalogue, the remaining mappable genome regions were assigned to background, and tiled at 5kb windows. See Supplemental Note for more detailed procedures.

We quantified the number of reads covering each region in the catalogue in each of our samples and atlas samples. We estimated a locally adaptive model of non-specific reads along the genome for each of the samples, and extracted counts that represent specific ChIP signal in the catalogue for each sample (Supplemental Note). These were then normalized (Supplemental Note) and scaled to 1M reads in the reference healthy samples.

#### Estimating capture rates

To estimate capture rates of cfChIP-seq we use two different approaches (see Supplemental Note for more details).

In the **global approach**, we compare input to output of the cfChIP-seq assay (Figure S4B). At the input end, we estimate the total number of nucleosomes that are present in the sample using the input cfDNA, which provides an upper bound on the number of nucleosomes it can contain (with each nucleosome ∼ 200bp of DNA). We also estimate the percent of these that are modified, which for H3K4me3 tend to be ∼1-2%. At the output end, we estimate how many of the unique fragments are background and how many are signal (see above). We then divide #signal fragments in output by #modified nucleosomes in input to get specific capture rate, and similarly #background fragment in output by total # nucleosomes to get non-specific capture rate.

In the **local approach**, we compare expected input coverage to output coverage (Figure S4C). Using input cfDNA amounts we can estimate the number of alleles (genomes) that cover each position. We then examine two types of regions, one “high-signal” where we assume that ∼100% of the nucleosomes are modified (e.g., promoters of constitutive genes) and the other one as “no-signal” where 0% of the nucleosomes are modified (e.g., background regions). The coverage we observe in the cfChIP-seq output is due only to non-specific capture in the no-signal region, and due to both specific and non-specific capture in the high-signal region.

#### Tissue Signatures

To define tissue specific signatures of a specific modification, we examined binned representation of the atlas according to our catalogue. For each tissue we defined a signature of unique windows with signal in one of the samples of the target tissue and without coverage in all others (Supplemental Note).

#### Gene level analysis

For each gene we defined the set of windows that match the gene (TSS in H3K4me3/2 and gene body in H3K36me3). The signal for a gene is the aggregate signal-background over windows associated with it (Supplemental Note).

#### Comparison to RNA-seq

The comparison of H3K4me3 ChIP to RNA-seq was performed as follows. RNA expression (normalized TPM) was downloaded from Roadmap Epigenomics Project (Table S10). Normalized cfChIP-seq coverage per gene in the matching sample was taken from the Roadmap Epigenomics Atlas (above). We examined RefSeq genes that appeared in both datasets. For each gene we computed pearson correlation between log(TPM+1) and log(ChIP-seq coverage+1) values across all ^56^ tissue/cell types that had match RNA-seq and H3K4me3 ChIP-seq data.

#### Estimating mean and variance

To define the healthy reference of signal per gene, we estimated the mean and variance of each gene in a set of 26 reference samples. The observed variation among the samples is due to the combination of biological variability and sampling noise. Thus, to estimate mean/variance we used a maximum likelihood approach that models the sampling noise of each sample and identifies the mean/variance that best matches this model (Supplemental Note).

#### Statistical analysis

We test whether a signature is present in the analysis of Figures 4 and 5. Formally, we examined whether we can reject the null hypothesis that the number of reads in signature windows is Poisson distributed according to background rate (Supplemental Note). We compute the p-value of the actual number of observed reads in signature windows as the probability of having this number or higher according to the null hypothesis. Rejection of the null hypothesis for a specific signature is an indication that some of the windows in the signature carry the modification in question in a subpopulation of cells contributing to the cf-nucleosome pool.

The second test is whether a gene presents a high signal with respect to its level in the baseline of healthy donors (Figure 5G). We use the signal from ^26^ healthy samples (Table S1) to estimate the mean and variance of reads in each region of interest (e.g., gene promoter). We then estimate two sample-specific parameters: 1) background rate (discussed above) and 2) a scaling factor that rescales average expectations to the sequencing depth of the specific sample (Supplemental Note). Together, these define the expected coverage of each gene-associated group of windows under the null-hypothesis that the subject is from the healthy population. We compute the p-value of the actual number of observed reads in the gene windows as the probability of having this number or higher according to the null hypothesis.

#### Pathways, and Transcription Factor targets

We downloaded a large collection of gene expression signatures representing different cellular processes, protein complexes, and transcriptional responses from the MSigDB collection ^53^. We downloaded transcription factor targets from Harmonizome database ^80^. These include targets from ENCODE ^81^, TRANSFAC ^82^, and CHEA ^83^.

#### Estimation of liver percentage

We used a linear regression model that matches the observed counts of a select representative genes to a sum of contribution of healthy-wo-liver and healthy liver. Briefly, we use the Roadmap Epigenomics Atlas “liver” (E066) as 100% liver. We assume that the mean healthy profile contains about ∼3% liver contribution, and so define the healthy-wo-liver as the result of subtracting 3% of liver profile from the healthy sample. We then identify the set of genes that are high in one of the profiles and close to 0 in the other. These are used as input features for robust linear regression (R rlm() function) that estimate the linear combination of liver and healthy-wo-liver profiles that is closest to the observed profile. The weights (linear regression coefficients) are normalized to sum to one, and the contribution of liver is taken as % liver in the sample.

#### Cancer signatures

We tested a compendium of gene programs from multiple sources (Table S10) against high-scoring CRC samples. Gene programs that had significant enrichment above/below healthy reference in at least 3 CRC samples but less than ⅔ of all the CRC samples were selected for the next step. The pattern of significantly above/below enrichments were clustered (Figure S7E). Each cluster of gene programs corresponds to a classification of the CRC samples (significant vs non-significant). For each such cluster we identified the genes that have significantly higher signal in the positive class of CRC samples compared to remaining CRC samples. The differential genes define a new gene-signature. These were clustered based on their classifications of samples, and combined into non-overlapping set of gene signatures (Supplemental Note).

### Data and Code availability

Code for processing cfChIP data is at https://github.com/nirfriedman/cfChIP-seq.git. Raw sequencing data was deposited to the EGA (EMBL-EBI) repository. BED files and browser tracks can be accessed at https:/Idoi.org/10.5281/zenodo.3967254.

