## Supplemental Note for "ChIP-seq of plasma cell-free nucleosomes identifies cell-of-origin gene expression programs"

### Supplemental Figures, Table captions, and Note

**Supplemental Figures, Table captions, and Note** 1
- Supplemental Figure S1 2
- Supplemental Figure S2 5
- Supplemental Figure S3 6
- Supplemental Figure S4 8
- Supplemental Figure S5 9
- Supplemental Figure S6 11

**Supplemental Tables** 14

**Supplementary Note** 15
- Assay reproducibility 16
- TSS location catalogue 19
- Enhancer location catalogue 20
- Gene body location catalogue 20
- Processing of sequencing files 21
- Estimating background signal 21
- Gene-level signal and normalization 23
- Estimate of cfChIP-seq capture efficiency 24
- Estimation of sequencing efficiency 28
- Defining tissue-specific signatures 31
- Statistical tests 32
- Cancer programs 35
- References 36

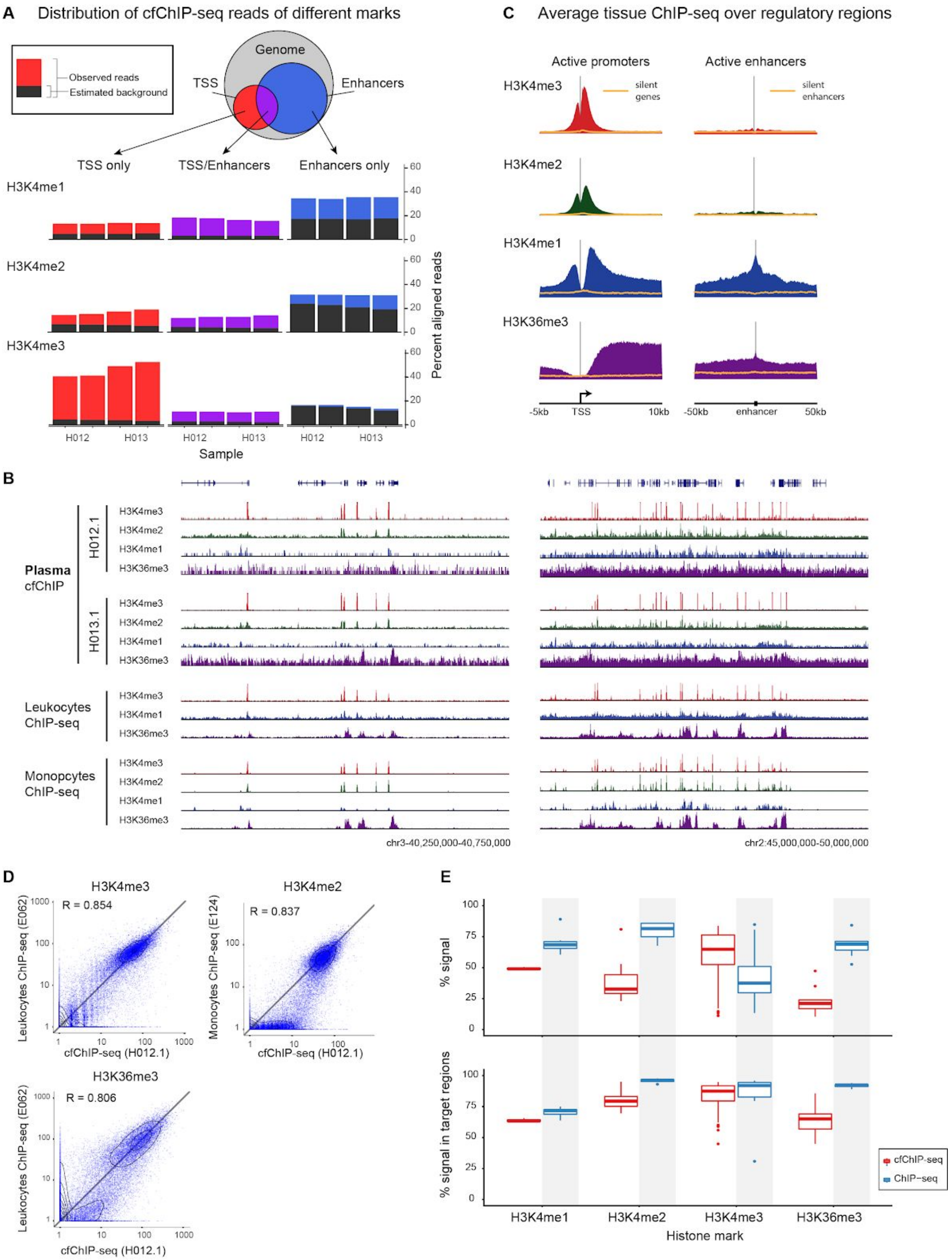

A Distribution of cfChIP-seq reads of different marks
Observed reads
Estimated background
Genome
TSS
Enhancers
TSS only
TSS/Enhancers
Enhancers only
H3K4me1
H3K4me2
H3K4me3
Percent aligned reads
H012
H013
Sample
C Average tissue ChIP-seq over regulatory regions
Active promoters
Active enhancers
silent genes
silent enhancers
H3K36me3
-5kb
TSS
10kb
-50kb
enhancer
50kb
B
Plasma cfChIP
H012.1
H013.1
Leukocytes ChIP-seq
Monopcytes ChIP-seq
chr3-40,250,000-40,750,000
chr2:45,000,000-50,000,000
D
R = 0.854
R = 0.837
R = 0.806
Leukocytes ChIP-seq (E062)
Monocytes ChIP-seq (E124)
cfChIP-seq (H012.1)
E
% signal
% signal in target regions
cfChIP-seq
ChIP-seq
Histone mark

#### Supplemental Figure S1

A. Distribution of reads for cfChIP-seq with different antibodies on four samples (H012.1, H012.2, H013.1, and H013.2). We divided the genome into regions that contain (putative) TSS based on our catalogue (see below) and (putative) Enhancers. Since there are regions that are marked as both (in different tissues), we consider the intersection separately. For each subset we show the fraction of reads mapped to the region. Within each bar, the fraction estimated as background (based on our background model, Methods) is marked in dark gray.
B. Genome browser view (as in Figure 1C).
C. Metaplots (as in Figure 1D) of ChIP-seq samples from the Roadmap Epigenomics compendium (1).
D. Scatter plots showing signal from cfChIP-seq versus Leukocyte ChIP-seq of H3K4me3, H3K4me2, and H3K36me3 (similar to Figure 1E).
E. Estimation of the amount of specific reads in cfChIP-seq. Top panel: box plot of the estimate of %reads that are above background levels for all the cfChIP-seq samples analyzed in the manuscript (Table S1) compared to selected ChIP-seq samples from Roadmap Epigenomics compendium (1). Bottom panel: percent of the signal above background that is in the expected genomic locations (i.e H3K4me1 and H3K4me2 - promoters and enhancers, H3K4me3 - promoters, H3K36me3 - gene bodies). For comparison, the same analysis pipeline was applied to selected Roadmap Epigenomic ChIP-seq samples against the same marks.

**A** Plasma cfChIP Fragment length

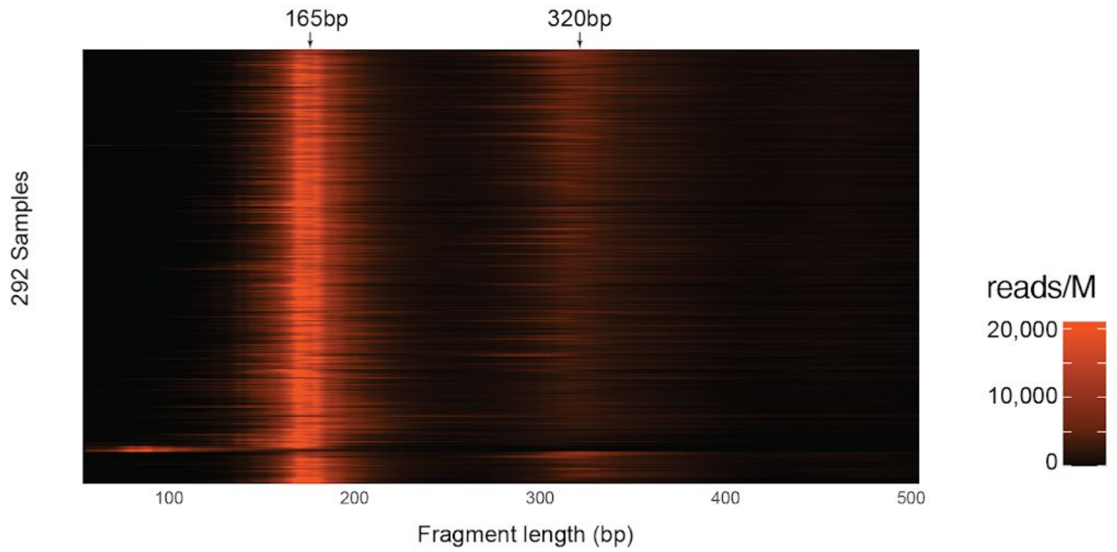


**B** Comparison of normalized gene counts

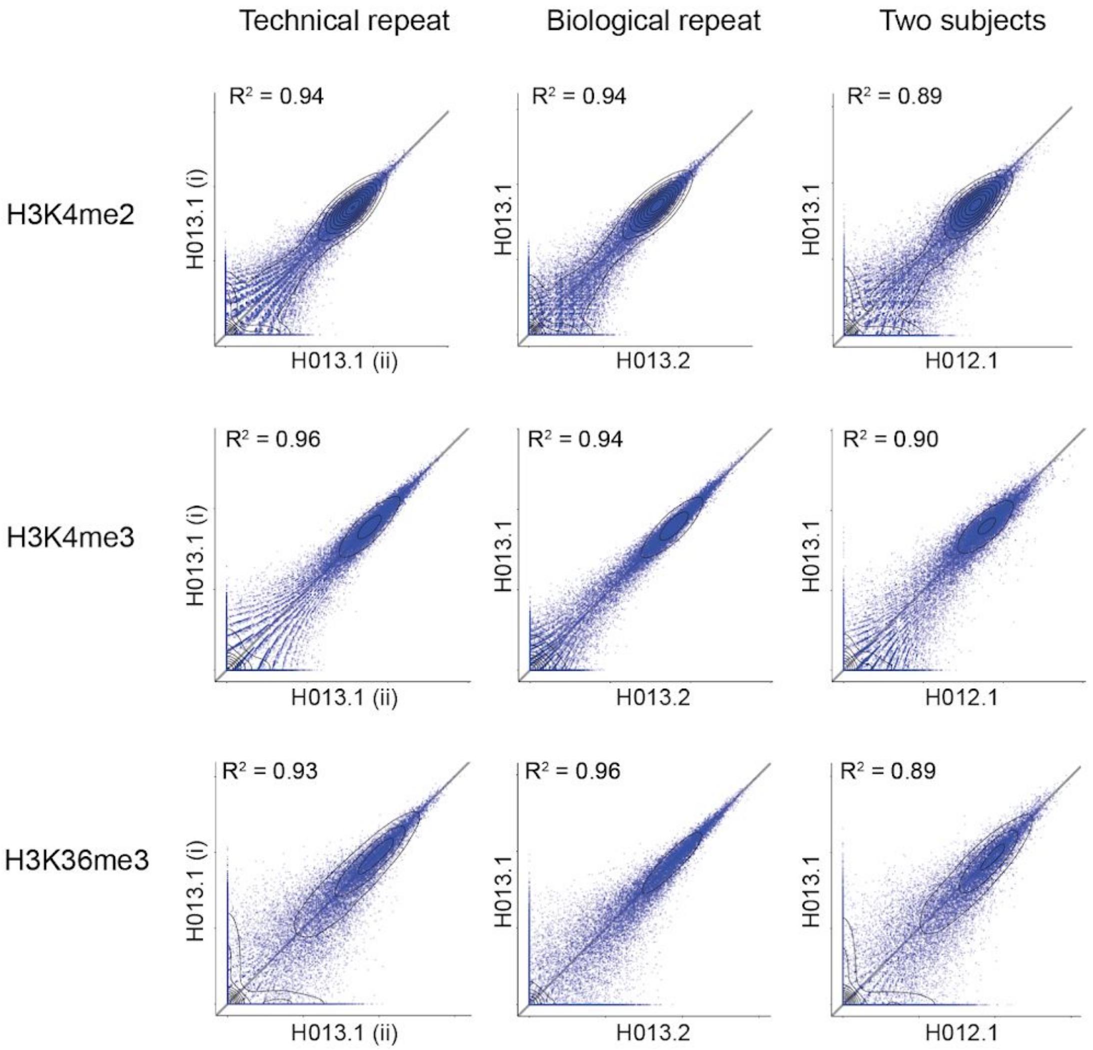

#### Supplemental Figure S2

A. Fragment length distribution for all samples in this manuscript. Each row represents a histogram of fragment length of a specific sample (RPM).

B. Reproducibility of the cfChIP-seq assay. Shown are technical repeats, biological repeats (two samples from the same donor) and comparison of two different donors for three histone marks. Each dot is a gene, and values are normalized counts at the gene promoter (H3K4me2/3) or body (H3K36me3).

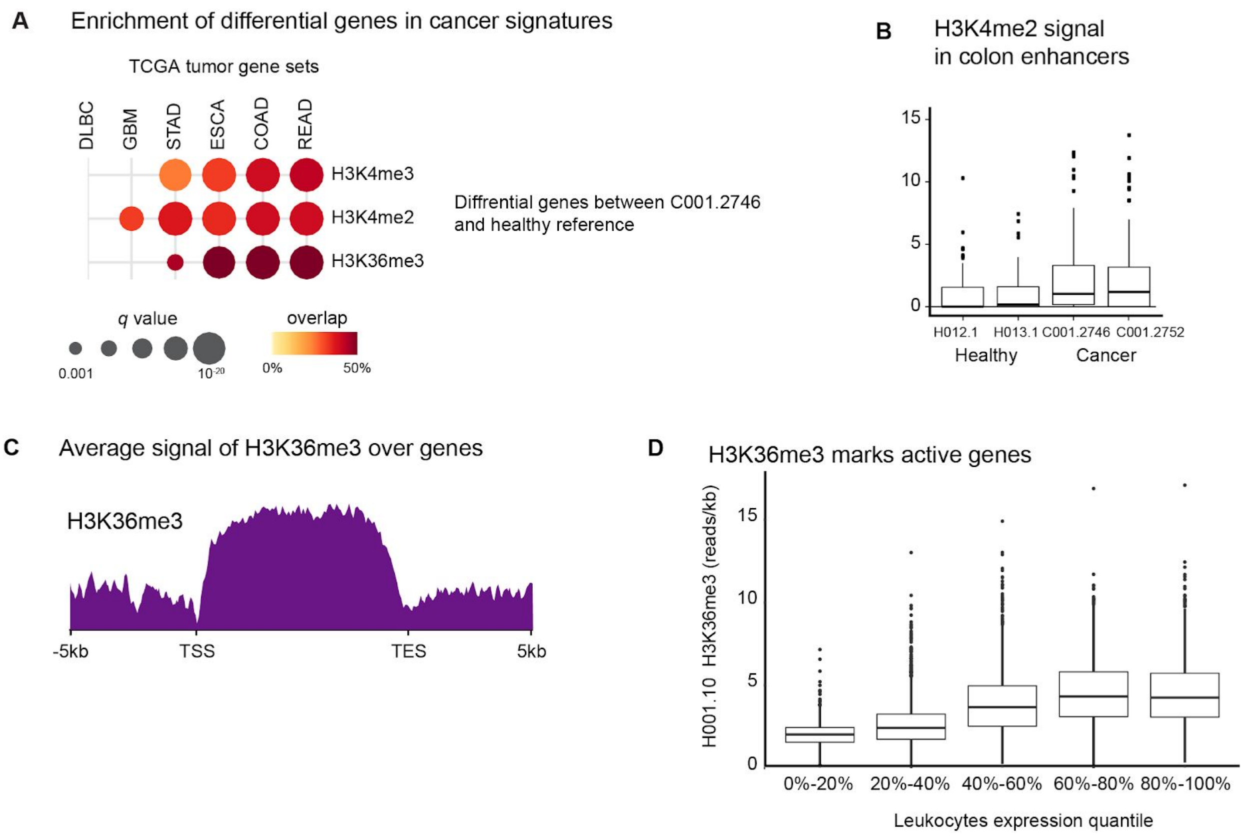


#### Supplemental Figure S3

A. Testing gene sets defined by highly expressed in different cancer types (TCGA, Methods) against genes with higher signal in a CRC tumor sample (Figure 2A).
B. Levels of H3K4me2 coverage over colon-specific enhancers in healthy donors and in CRC cancer samples.
C. Average coverage of H3K36me3 across gene bodies (meta gene)
D. Average coverage of H3K36me3 over gene bodies for genes at different expression quantiles.

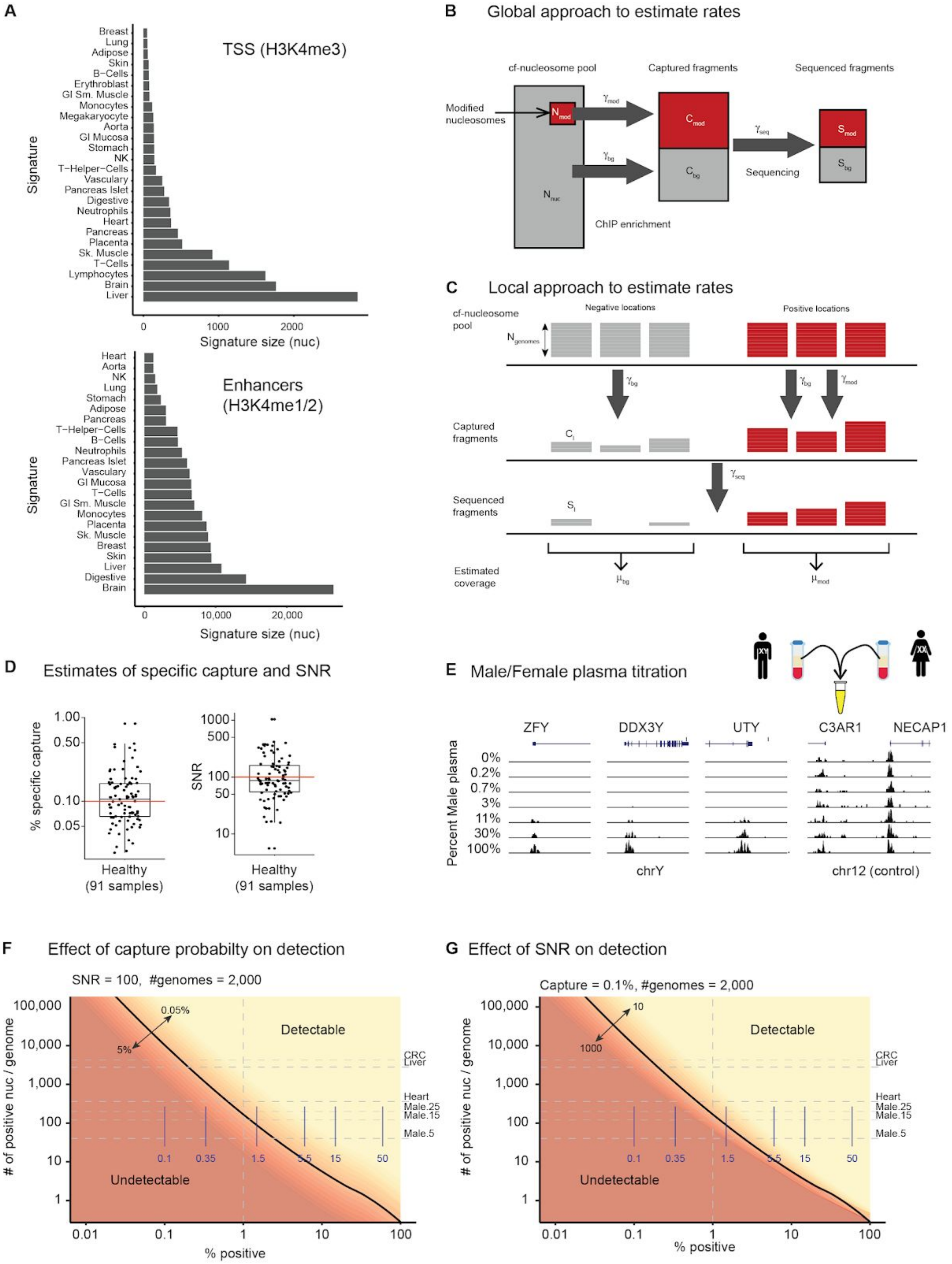
A
TSS (H3K4me3)
Breast
Lung
Adipose
Skin
B−Cells
Erythroblast
GI Sm. Muscle
Monocytes
Megakaryocyte
Aorta
GI Mucosa
Stomach
NK
T−Helper−Cells
Vascuiary
Pancreas Islet
Digestive
Neutrophils
Heart
Pancreas
Placenta
Sk. Muscle
T−Cells
Lymphocytes
Brain
Liver
Signature
0
1000
2000
Signature size (nuc)
Enhancers
(H3K4me1/2)
Heart
Aorta
NK
Lung
Stomach
Adipose
Pancreas
T−Helper−Cells
B−Cells
Neutrophils
Pancreas Islet
Vascuiary
GI Mucosa
T−Cells
GI Sm. Muscle
Monocytes
Placenta
Sk. Muscle
Breast
Skin
Liver
Digestive
Brain
Signature
0
10,000
20,000
Signature size (nuc)
B
Global approach to estimate rates
cf-nucleosome pool
Captured fragments
Sequenced fragments
Modified nucleosomes
N mod
γ mod
C mod
γ seq
S mod
γ bg
C bg
Sequencing
S bg
N nuc
ChIP enrichment
C
Local approach to estimate rates
cf-nucleosome pool
Negative locations
Positive locations
N genomes
γ bg
γ bg
γ mod
Captured fragments
C i
γ seq
Sequenced fragments
S i
Estimated coverage
μ bg
μ mod
D
Estimates of specific capture and SNR
% specific capture
1.00
0.50
0.10
0.05
Healthy
(91 samples)
SNR
1000
500
100
50
10
Healthy
(91 samples)
E
Male/Female plasma titration
ZFY
DDX3Y
UTY
C3AR1
NECAP1
Percent Male plasma
0%
0.2%
0.7%
3%
11%
30%
100%
chrY
chr12 (control)
F
Effect of capture probabilty on detection
SNR = 100, #genomes = 2,000
### of positive nuc / genome
100,000
10,000
1,000
100
10
1
0.05%
5%
Detectable
Undetectable
CRC
Liver
Heart
Male.25
Male.15
Male.5
0.1
0.35
1.5
5.5
15
50
0.01
0.1
1
10
100
% positive
G
Effect of SNR on detection
Capture = 0.1%, #genomes = 2,000
### of positive nuc / genome
100,000
10,000
1,000
100
10
1
10
1000
Detectable
Undetectable
CRC
Liver
Heart
Male.25
Male.15
Male.5
0.1
0.35
1.5
5.5
15
50
0.01
0.1
1
10
100
% positive

#### Supplemental Figure S4

A. Total sizes (in nucleosomes) of TSS (Top) and Enhancer (Bottom) signatures of various cell types.

B. Global approach to estimate capture rates. Based on the known fraction of the genome modified and the number of input genomes (based on ng/ml measurements) we estimate the number of modified and unmodified nucleosomes in the cfChIP-seq sample. We estimate the fraction of reads that are signal (specific capture) and background (non-specific capture) in the sample. The ratio between these is an estimate of capture probability * probability of sequencing captured fragments. Dividing by the estimate of sequencing efficiency (Supplementary Note), we recover capture rates.

C. Local approach to estimate capture rates. Using the number of input genomes, we assume that in high house-keeping locations all input nucleosomes are modified while in regions without promoters none of the input nucleosomes are modified. Computing the coverage over these two types of regions we recover specific and non-specific capture rates.

D. Estimates of specific capture rate and of SNR (specific capture / non-specific capture) over 88 healthy samples, assuming 1000genomes/ml and 2ml input.

E. Genome browser of chrY male-specific promoters (left) and a representative autosomal region (right) in the male/female titration experiment.

F. Simulation study of the effect of capture probability on detection. The blue marks denote the concentrations used in the male-female titration experiment which had capture probabilities ~0.1% and SNRs of ~500-800 (Figure 3D).

G. Simulation study of the effect of SNR levels on detection probability.

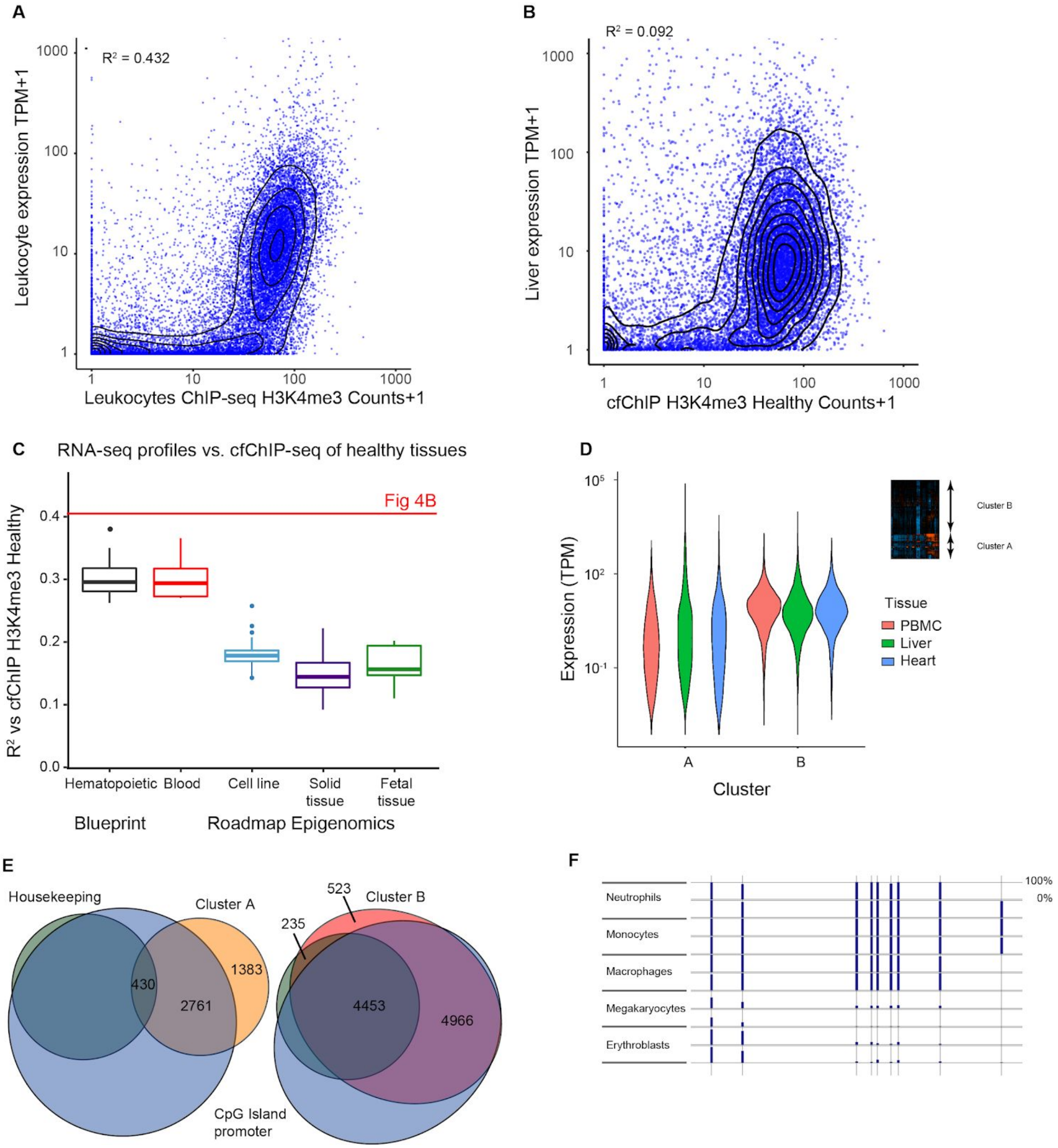


#### Supplemental Figure S5

A. Comparison (as in Figure 4B) of Leukocytes H3K4me3 ChIP-seq signal vs. Leukocytes gene expression levels (both for Roadmap Epigenomic sample E062).

B. Comparison (as in Figure 4B) of H3K4me3 cfChIP-seq signal from a healthy donor (H012.1) vs. Liver gene expression levels (Roadmap Epigenomics sample E066).

C. Summary of correlations of healthy cfChIP-seq levels against different expression patterns from Roadmap Epigenomics and BLUEPRINT. For each category of expression profiles we plot the boxplot of $R^2$ values.

D. Comparison of the expression levels of genes in two clusters of Figure 4C (see inset). Cluster A contains 4,690 genes that change between samples, and Cluster B contains 10,177 genes that do not change between samples. Violin plots show the distribution of expression levels in three tissues - PBMC, Heart, and Liver, from the Roadmap Epigenomics expression data.

E. Overlap of both clusters with the set of genes with CpG island promoters (blue) and housekeeping genes (green; based on analysis of GTEX compendium, see Methods). For clarity we show each cluster in a separate Venn diagram.

F. Percent of methylation levels at CpGs in the FECH intron, as examined in Figure 1 of Lam et al (2) in samples of several cell types in the BLUEPRINT project**.** Shown are two samples from each cell type. The blue bars represent the level of CpG methylation detected at each of the CpGs (marked by vertical gray lines). Missing estimates are due to low coverage.

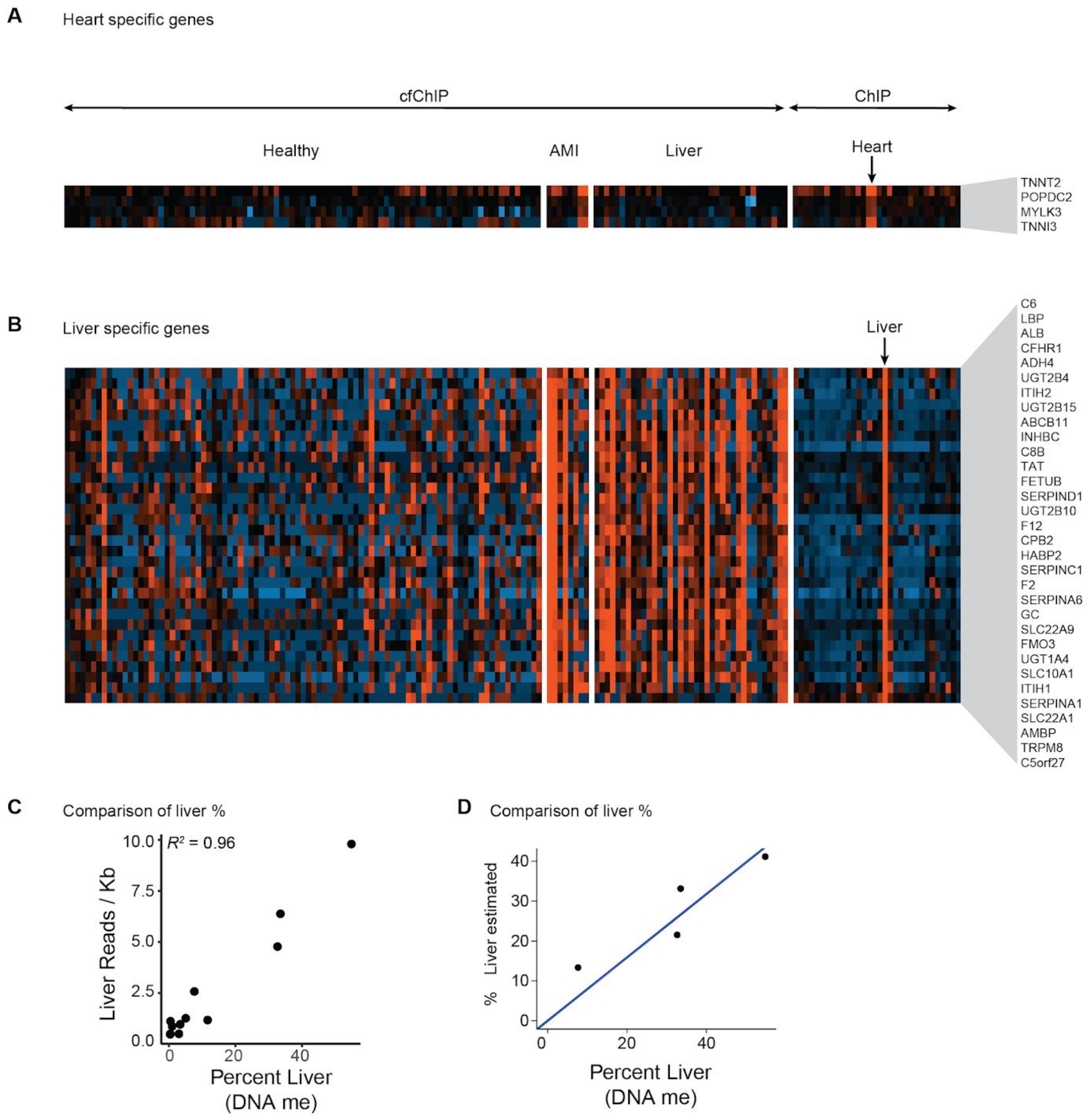


#### Supplemental Figure S6

A. Zoom in on the heart specific genes in Figure 4D.

B. Zoom in on the liver specific genes in Figure 4D.

C. %Liver DNA Me vs Signature strength

D. %Liver DNA Me vs estimate of % liver (cfChIP-seq)

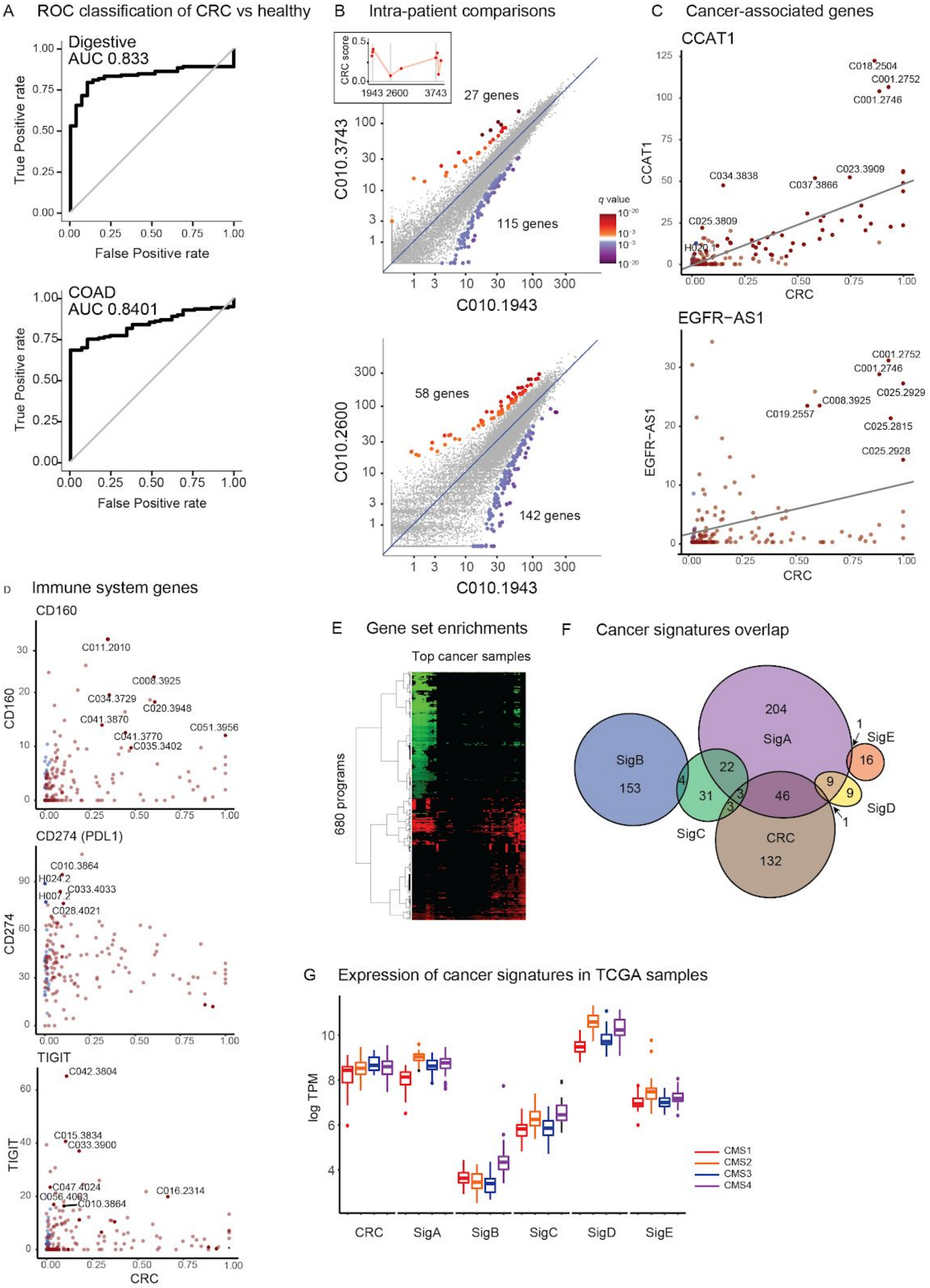

A ROC classification of CRC vs healthy
Digestive AUC 0.833
COAD AUC 0.8401
True Positive rate
False Positive rate
B Intra-patient comparisons
CRC score
1943 2600 3743
27 genes
115 genes
C010.3743
C010.1943
58 genes
142 genes
C010.2600
q value
C Cancer-associated genes
CCAT1
C018.2504
C001.2752
C001.2746
C034.3838
C037.3866
C023.3909
C025.3809
H020.1
CRC
EGFR−AS1
C001.2752
C001.2746
C025.2929
C008.3925
C019.2557
C025.2815
C025.2928
D Immune system genes
CD160
C011.2010
C008.3925
C034.3729
C020.3948
C041.3870
C051.3956
C041.3770
C035.3402
CD274 (PDL1)
C010.3864
H024.2
C033.4033
H007.2
C028.4021
CD274
TIGIT
C042.3804
C015.3834
C033.3900
C047.4024
C016.2314
C056.4003
C010.3864
E Gene set enrichments
Top cancer samples
680 programs
F Cancer signatures overlap
SigB 153
204 SigA
SigE 16
22
4
31
3
3
46
9
9
1
1
SigD
SigC
CRC 132
G Expression of cancer signatures in TCGA samples
log TPM
CMS1
CMS2
CMS3
CMS4
CRC
SigA
SigB
SigC
SigD
SigE

#### Supplemental Figure S7

A. Performance of CRC samples classification with Digestive (Top) and COAD (Bottom) signatures (as Figure 7C).

B. Intra-patient comparisons (as Figure 7E). Inset: time samples drawn on the patient timeline (Figure 7D)

C. Levels of CRC associated genes in different samples. Each point is a sample (red - cancer, blue - healthy) plotted with %CRC (x-axis) vs normalized number of reads of the gene (y-axis). Solid points - the signal of the gene is significantly above background.

D. Example of immune-related genes in CRC samples. Same as (similar top (C) )

E. Clustering of gene set enrichment in CRC samples (see Table S8).

F. Venn diagram of overlaps between cancer signatures.

G. Evaluation of cancer signatures in CRC samples from TCGA, grouped by their CMS subtype.

### Supplemental Tables

**Table S1: Sequencing statistics for each sample**

Sequencing statistics of the samples sequenced in this study.

**Table S2: Tumor signatures**

List of significant genes in each of the TCGA cancer types (Methods). Each line starts with the cancer type name followed by a list of genes.

**Table S3: Subjects and sample clinical information**

For each subject we list date of birth and relevant clinical details. For each sample we list the date it was collected. Tabs separated by Healthy, AMI, Liver, and CRC.

**Table S4: Cell type signatures**

Each line lists one genomic window with the following fields: Signature: name of the signature (e.g., Liver, or T-cells), Chr: chromosome, Start: start location, End: end location, Gene: gene name if there is a gene (or genes) associated with the location, Type: TSS or background., Browser: browser address line that contains the region + 10kb to each direction, Signal: maximal normalized signal in this window in tissues that belong to the positive class, Background: maximal normalize signal in tissues that do not belong to the positive class.

**Table S5: Full analysis of tissue signatures vs samples**

Each line lists one sample vs. one signature. Columns: Sample: sample name; Signature: signature name; Observed counts: number of reads in the signature; Background: total estimate of background reads in the signature; Normalized counts: normalized counts; p-value: Poisson significance test above background (-log10 of p-value); q-value: FDR correction per sample (-log10 of q value); Z-score - difference in actual reads from expected by standard deviation.

**Table S6: Full analysis of gene sets vs samples with reference**

Each line lists one sample vs. one gene set. Columns: Sample: sample name; Gene set: gene set name from MSigDB curated collection (3); Observed counts: number of reads in the signature; Background: total estimate of background reads in the signature; Normalized counts: normalized counts; Normalized expected: the expected level in healthy donors; p-value: two-tailed test(-log10 of p-value); p-value above/below: single tailed test (-log10); q-value: FDR correction per sample (-log10 of q value); Z-score -

difference in actual reads from expected by standard deviation.

**Table S7: Liver - Cluster enrichments**

Each line is a test of a cluster against one term as performed by EnrichR (4). Columns: Cluster: cluster name; DB: database name; Term: name of term; Query size: size of cluster; Overlap: overlap of query with term; p-value: hypergeometric p-value; Adjusted p-value: adjusted p-value; Genes: names of genes in the overlap.

**Table S8: Gene set counts in CRC samples relative to healthy reference**

Each row represents the result for a single gene set. The columns represent CRC samples with relatively high cancer load (CRC Signature > 0.15). The values shown are the gene set counts relative to healthy reference. Only significant results are shown.

**Table S9: CRC - Signature enrichments**

Each line is a test of a signature against one term as performed by EnrichR (4). Columns: Signature: signature name; DB: database name; Term: name of term; Query size: size of cluster; Overlap: overlap of query with term; p-value: hypergeometric p-value; Adjusted p-value: adjusted p-value; Genes: names of genes in the overlap.

**Table S10: Data sources**

**Table S11: Roadmap samples used**

### Supplementary Note

#### Assay reproducibility

To assess the reproducibility of cfChIP, we performed technical and biological replicates from several subjects. Specifically, two healthy individuals (H012 and H013, one female and one male) were sampled in two biological repeats, after finalizing the development of the experimental and analysis methods. These samples were not part of our method development as objective test cases. This experiment was run once and we used the two samples without any cherry picking.

Reproducibility can be assayed at multiple levels. The simplest is to compare the raw cfChIP signal of samples. We tiled the genome with 2kb tiles, and counted the number of reads mapped onto each tile. As might be expected, most of the tiles had zero observations, but some had higher values. See Figure S8 for some examples.

This comparison shows that some samples have differences in yields (either ChIP or sequencing efficiency, see below). In general the $R^2$ values for H3K4me3 are high (~0.8-0.95), for H3K4me2 are somewhat lower (~0.6-0.8) and much lower for H3K4me1 and H3K36me3. This is consistent with the sharply concentrated nature of H3K4me3 peaks, and the much more diffuse patterns of H3K4me1 and H3K36me3.

A more relevant comparison of samples is by examining the estimated counts per gene. These estimates remove non-specific background and normalize samples to reduce artifacts of differential yield levels. Indeed, comparing normalized gene signal profiles shows much better agreement between healthy samples (Figure S2B).

When examining all 88 healthy samples of H3K4me3, we can see that normalized gene correlations are much better than raw 2kb counts (compare Figure S9A vs Figure S9B). In particular biological repeats, tend to be of higher correlations.

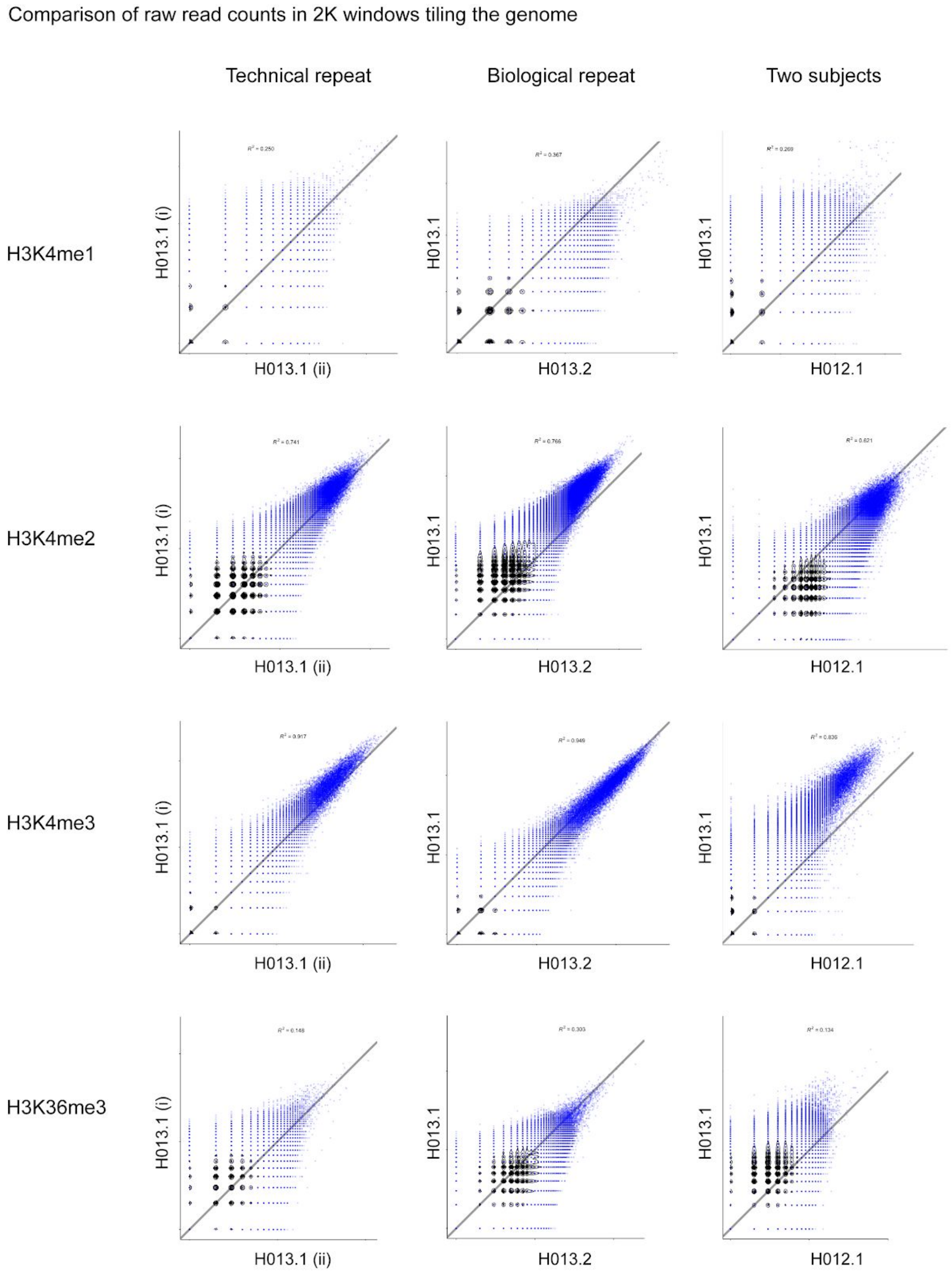


**Figure S8: Comparison of signal in 2kb tiles along the genome.**

For each modification (rows) we compared two technical repeats (left column), biological repeats (middle column) and two donors (right column). Each plot shows log(1+count) in one sample to the other. Gray line indicates the x=y line.

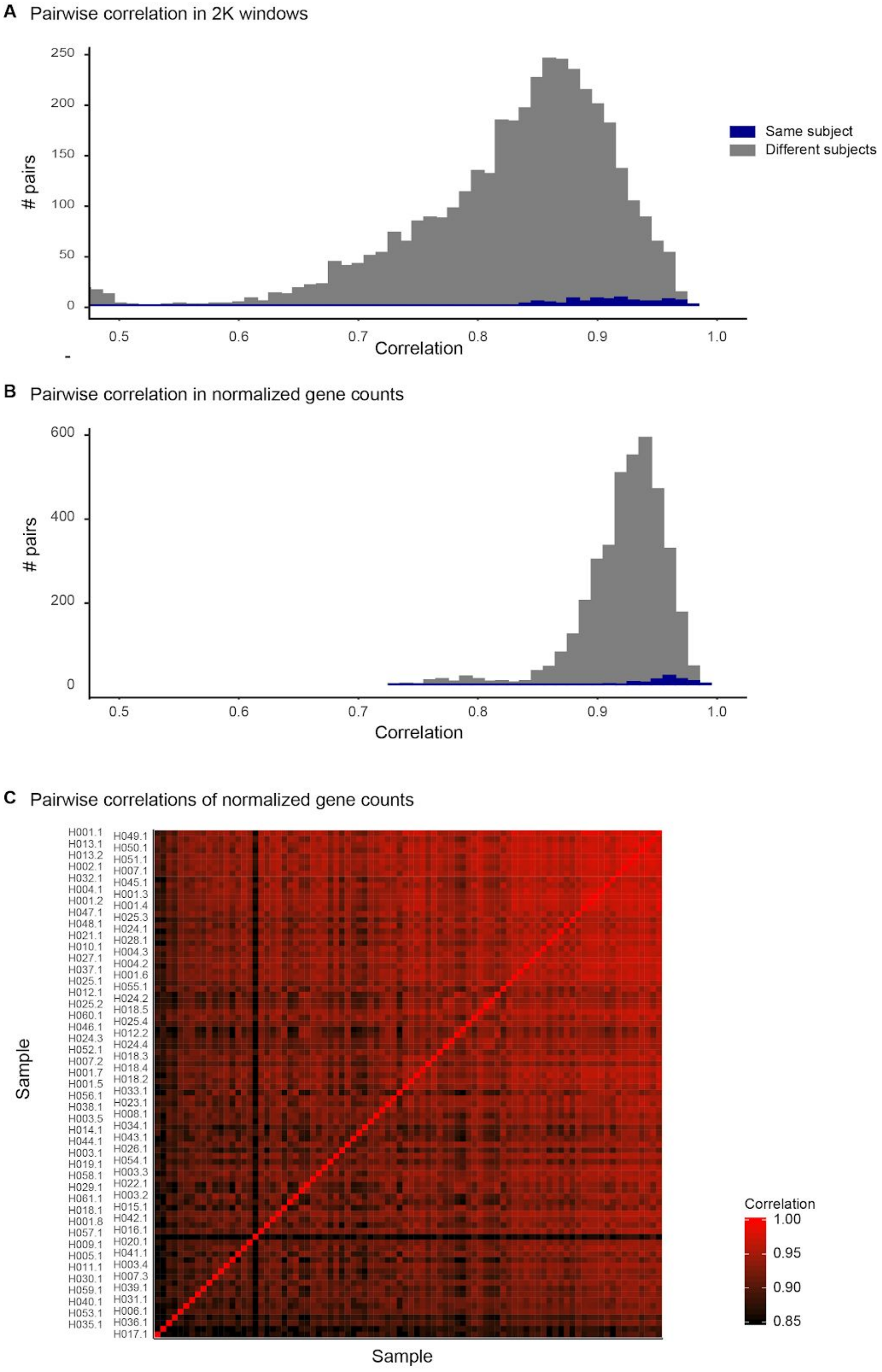


**Figure S9: Reproducibility of H3K4me3 profiles.**

A. Histogram of pairwise correlations between healthy samples at the level of 2kb tiles.
B. Histogram of pairwise correlations between normalized gene counts.
C. Heatmap showing pairwise correlations of gene counts.

#### TSS location catalogue

We constructed a TSS catalogue through the following steps. All steps were carried on the "hg19" human genome assembly:

1. We downloaded ChromHMM calls for 111 tissues and cell types throughout the human genome from the Roadmap Epigenomics website (Table S10). UCSC Browser known gene annotations and ENSEMBL transcript annotations were downloaded (Table S10).

2. We filtered all genomic ranges that were marked with states "1_TssA" or "2_TssAFlnk", and merged adjacent ranges that were marked as either state in exactly the same set of tissues. We call these "ChromHMM TSS windows". We found 640,744 such windows. Each ChromHMM TSS window was assigned gene name(s) through the following steps:

    a. If it was within 2.5Kb of one or more TSS in UCSC known gene annotation, it was assigned the name of these genes.

    b. If not, we searched for an ENSEMBL transcript starts within 2.5Kb. Again if such were found the TSS window received the gene name associated with the transcript.

    c. All other TSS windows remained without a name

    This procedure assigned names to 190,552 windows, and 450,192 windows remained unnamed. In some cases the unnamed TSS windows can be seen as alternative starts of an annotated gene (either upstream or downstream of the annotated TSS), but in many other cases these were far from transcript starts. In general, after correcting for length, the rate of seeing a read from H3K4me3 cfChIP in unnamed windows is 6-fold smaller than that of named windows. This suggests that most of the unnamed windows are not observed in our data. In terms of coverage, named TSS windows cover 4.46% of the genome, and unnamed ones an additional 7.03%. Given the estimate of 1-2% of nucleosomes in a cell carrying the H3K4me3 mark, we conclude that most of these putative promoters are not used in cell types we observe. Given observations of H3K4me3 signal at enhancers (5), these might be signatures of active enhancers, but this would require additional verification.

3. To include transcripts that are not represented in the TSS catalogue, we examined all genes in the UCSC known gene database and all transcripts in the ENSEMBL database. For each gene we defined a TSS window of size 3Kb centered on the TSS. We discarded all such windows that overlapped with a TSS window described in step 2. In total this step added 11,518 and 36,166 TSS windows from UCSC known genes and ENSEMBL transcripts, respectively.

4. We created windows that tile the remaining genomic regions between TSS windows. For each TSS window without an adjacent TSS window, we created “flanking” regions of size 1Kb (or less in case of collisions). This resulted in 475,727 flanking windows (as some of the TSS windows are adjacent to each other, depending on ChromHMM calls in different tissues). The remaining uncovered regions were tiled with “background” regions of size 5Kb (or less). In total there were 485,245 such background windows.

The resulting catalogue was saved as a BED file (TSS.bed). This catalogue was used in all analysis of H3K4me3 cfChIP and re-analysis of reference ChIP-seq data.

#### Enhancer location catalogue

The construction of enhancer catalogue was similar to that of the TSS catalogue, except for Step 2 above where we included the enhancer states “6_EnhG” and “7_Enh”. Similarly to the TSS catalogue construction, we merged consecutive enhancer windows that are called in the same tissues. In Step 3 we only added additional TSSs (as above). Step 4 was left unchanged, except for the fact that we considered regions that were neither TSS nor Enhancers.

The resulting catalogue contained 2,345,831 enhancer windows, most of which were relatively short: 91.9% of the windows are 1kb or shorter, and 49.2% are only 200bp long (the unit length of ChromHMM calls). Nonetheless, these windows cover 40% of the genome.

We used this for the analysis of H3K4me1 and H3K4me2 cfChIP.

#### Gene body location catalogue

The construction of this catalogue was similar to that of the TSS catalogue, except that in Step 2 we collected only the transcribed states “3_TxFlnk”, “4_Tx”, and “5_TxWk”. We named regions if they overlapped with an annotated gene or transcript.

In Step 3 we added the UCSC known gene database and all transcripts in the ENSEMBL database and added the gene body, if it did not overlap with regions from Step 2. The resulting gene regions are often very long, and thus before Step 4, all gene regions were tiled with 5kb size windows. Step 4 was left unchanged, except it considered regions that were not genes.

The resulting catalogue contained 1,021,467 gene windows based on ChromHMM calls, and additional 4,301 and 861 gene windows based on UCSC genes and ENSEMBL transcripts, respectively, that were not covered by ChromHMM. These numbers are consistent with the fact that most genes were expressed in one of the tissues covered by the Roadmap Epigenomics project. Most of the genome (71%) was covered by these windows --- recall that these include both exonic and intronic regions.

We used this catalogue for the analysis of H3K36me3 cfChIP.

#### Processing of sequencing files

Base calling was performed with bcl2fastq (2.18/2.20). Paired-end reads were mapped to the human genomes (‘hg19’ assembly) using bowtie2 with “no-mixed” and “no-discordant” flags discarding reads with quality=0. BEDPE files (start and end of every fragment) were obtained using BEDtools “bamtobed” with “bedpe” flag, discarding duplicate fragments.

BEDPE files were converted to coverage counts over windows in the catalogue using BioConductor “GenomicRanges” countOverlaps() function. Specifically, we assigned a fragment to a window if its center is within the window. In other words, all fragments were resized to length 1 while maintaining their center.

#### Estimating background signal

Every ChIP procedure has a non-specific background signal. In the case of cfChIP the background is due to non-specific binding of DNA and chromatin fragments to the beads-antibody complex. Our experience showed that the background levels varied between samples and batches of bead-antibody ligation. Moreover, the sequencing depth varied between samples, and in a deeply sequenced sample the number of background reads increases. Thus, it was important to estimate background signal levels to be able to contrast them with actual signal.

We initially applied a simple minded procedure for removing background in H3K4me3 signal. We reasoned that virtually all of the specific signal in H3K4me3 is located in TSS and gene 5’ regions. Thus, reads in other locations represent background. To account for TSS that are not annotated in our TSS catalogue, we reasoned that some small fraction of background windows might contain real signal, and thus we ignored the ones with the highest values in estimation of background.

In more detail we did the following. We created a vector with the coverage of all “background” windows of size >= 4Kb (323,237 out of 485,245). The vast majority of these were 5Kb long. We then applied the following procedure:

estimateBackground $(X)$ // X vector of values

$T \leftarrow quantile(95, X)$ // find the 95th quantile of X

$X \leftarrow X[X \leq T]$ // restrict ourselves to values below T

$\hat{\lambda} = \arg\max_{\lambda} \prod_{i=1}^{|X|} P_{\lambda}(x_i \mid x_i \leq T)$ // maximum likelihood of truncated poisson

Return $\hat{\lambda}/5$ // convert to reads/Kb (the median length of // background windows is 5KB)

This procedure was relatively robust to the choice of quantile for removing outlier windows.

However, in some samples the Poisson distribution was not a good fit for background values. Further examination revealed that much of this discrepancy was due to local background effects. One example of local effect is the sex chromosomes that appear at 50% levels in male samples, and 100% (chrX) and 0% (chrY) in female samples. These were not the only local effects - some regions showed higher levels of background. This could be due to segmental duplications (mostly at regions close to centromeres and telomeres) or accessibility issues. Moreover, in cancer samples there were patient specific deviations that most likely reflect chromosomal aberrations.

To overcome these issues, we devised a localized background rate estimate. We repeated the estimation procedure described above, in different genomic resolutions:

1. Genome-wide background level.
2. Chromosome-specific background.
3. Tiles of 10Mb covering each chromosome at offsets of 2.5Mb.
4. Tiles of 5Mb covering each chromosome at offsets of 1.25Mb.

The estimate at each level used the estimate of the previous level as a prior (using pseudo-counts of 1000 windows in levels 2 and 3, and 500 windows in level 4).

The result is an estimate of background coverage rate at overlapping tiles of 5Mb. To get a single estimate, for each location we take the maximum of the estimate of the tiles covering it (typically 4 tiles). We chose the maximum as we reasoned that over-estimate of background might reduce the estimated signal but would minimize background artifacts.

Example of the background estimate for a healthy sample (male)

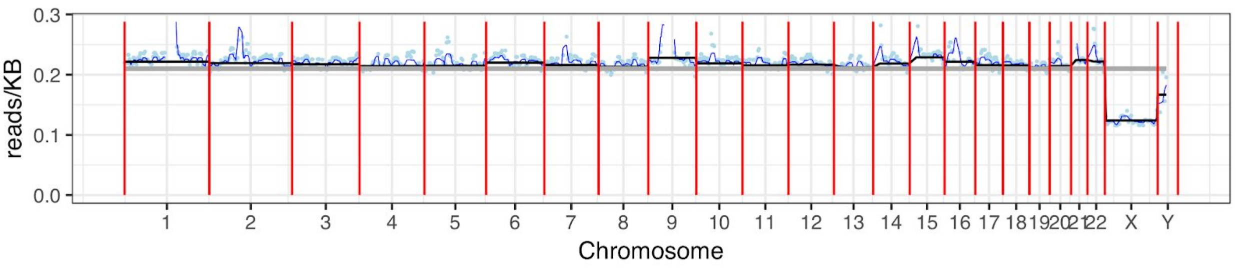


The gray line displays the genome-wide estimate, the black lines the chromosome-specific estimate, the blue line the larger tile-based estimates, and the light blue points the local estimate (based on the smaller tiles)

In this example the chrX background is lower than autosomal chromosomes (slightly more than half) and chrY background is a bit lower. Many locations in chrY are orthologous to ones in chrX leading to skewed estimates. Other deviations occur close to centromeres where we find the background level to be higher in some chromosomes (e.g., chr1, chr9).

A healthy female sample is similar, except for the sex chromosomes.

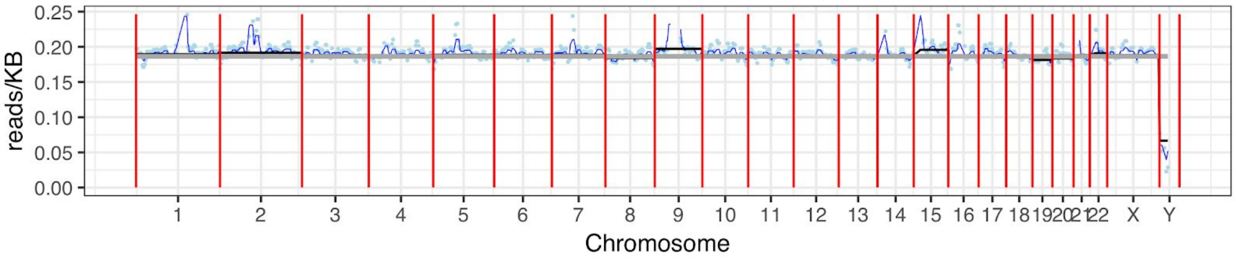


When we examine a patient with cancer, the background estimate is much more variable, presumably reflecting chromosomal aberrations in the tumor (e.g., duplication in chr13 and in an arm of chr8).

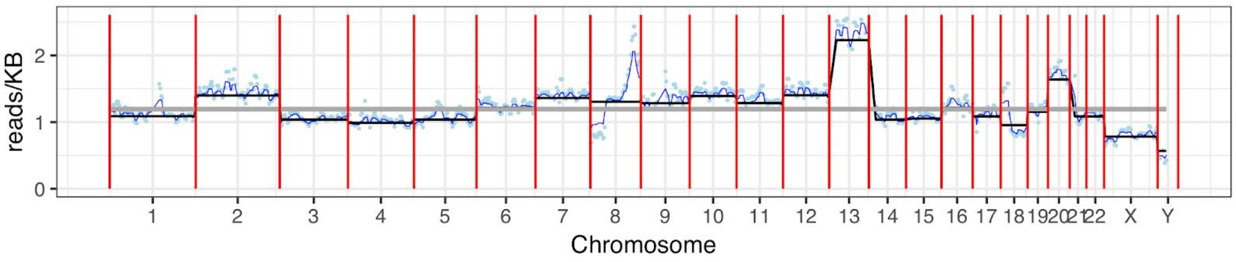


#### Gene-level signal and normalization

We define the following notation:

- $C[w, s]$ - read counts observed in window $w$ in sample $s$
- $\hat{\lambda}[w, s]$ - estimated background rate in window $w$ in sample $s$

Using these, we define

$$B[w, s] = \hat{\lambda}[w, s] * width(w)$$

To be the background rate for window $w$ in sample $s$. The rates we estimate are in units for counts/kb, and $width(w)$ returns the width of window $w$ in units of kb, and so the result is a rate in units of counts.

For each gene we assigned a set of TSS windows that are annotated with the gene name. For each sample we computed the actual total coverage over the windows assigned to the gene and the expected mean of background reads over these windows (taking into account the local rate and the width of each window).

More precisely,

$$C[g, s] = \sum_{w \in W_g} C[w, s]$$

$$B[g,s] = \sum_{w \in W_g} B[w,s]$$

where $W_g$ is the set of windows assigned to gene $g$.

The null assumption is that the coverage $C[g,s]$ is distributed as a Poisson with parameter $B[g,s]$. Thus, we argued that values much larger than expected are signals. We define the raw signal at gene $g$ as

$$S[g,s] = C[g,s] - B[g,s] \quad \text{if } C[g,s] \geq B[g,s] \quad \text{and 0 otherwise}$$

To normalize the effect of different coverages, we reasoned that the signal at promoters of "housekeeping" autosomal genes should be similar in different samples. We defined these genes as ones with highly significant signals in a set of reference healthy samples (Table S1). The precise choice of significance level did not change the normalization.

We applied quantile normalization (Bioconductor normalize.quantiles()) between the average signal of the housekeeping genes in the healthy reference set to the signal of the housekeeping genes in each one of the acquired cfChIP samples. This resulted in normalized values for housekeeping genes in each sample. However, it does not assign values for all other genes. We thus used linear regression to estimate a multiplicative normalization factor for each sample to best match quantile-normalized values to raw values. For most samples the relation between the two was linear.

The scaling factors were rescaled so that the total normalized signal (below) at the set of reference healthy samples will be a million on average.

Using these normalization factors, $\nu[s]$ we computed for each sample the normalized gene levels

$$N[g,s] = \nu[s] * S[g,s]$$

Using the same normalization procedure we also normalized the coverage at each window in each sample

$$N[w,s] = \nu[s] * \max(C[w,s] - B[w,s], 0)$$

#### Estimate of cfChIP-seq capture efficiency

To get a rough estimate of the cfChIP procedure efficiency, we consider two alternative estimates: a **global** method and a **local** method.

Both start by estimating the amount of material in the plasma.

Assuming one human diploid cell contains 6.6pg of DNA, we can use the measured amount of cfDNA in each sample to compute the number of genomes:

$$N_{genome} = 2 * V * C / 0.0066$$

Where $V$ is the sample volume (in ml), and $C$ is the cfDNA concentration (in ng/ml).

Assuming an average nucleosome consists of ~200bp (including linker), each genome is packed by ~ $1.65*10^7$ nucleosomes and we compute

$$N_{nuc} = N_{genome} * 1.65 * 10^7$$

This is likely an overestimate of the amount of input material, since not all cfDNA is necessarily packed in nucleosomes.

The **global method** is described in the following scheme:

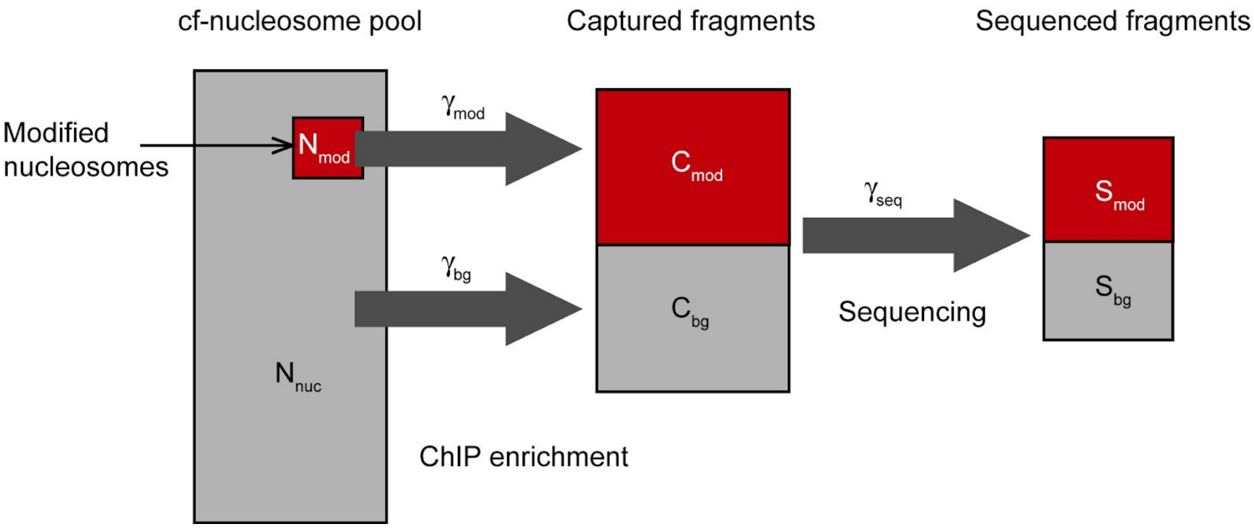


To estimate the total number of nucleosomes marked by a certain modification, we assume that the same percent of the genome is marked in every cell. Denote this to be ($p_{mod}$) we have:

$$N_{mod} = N_{nuc} * p_{mod}$$

If we estimate the number of marked nucleosomes we captured in total ($C_{mod}$) we can then estimate the yield as

$$\gamma_{mod} = C_{mod}/N_{mod}$$

If we also estimate the number of unmarked (background) nucleosomes we captured ($C_{bg}$) we can also estimate the rate of non-specific capture by comparing it to the total input

$$\gamma_{bg} = C_{bg}/N_{nuc}$$

This method raises two issues. 1) How do we estimate the percent of nucleosomes in a cell marked by every modification, and 2) how do we estimate the fraction of sequenced reads that originated from modified nucleosomes vs. background reads.

For the percent of marked nucleosomes we use a rough estimate based on ChIP-seq of specific blood cells (1) and other quantitative studies of modifications. Single molecule imaging (6) in multiple cell types show that H3K4me3 levels are fairly consistent at 2% while quantitative MS suggest less than 1% (7). Based on H3K4me3 ChIP-seq of blood cells we estimate the percent to be:

$$p_{K4me3} = 0.01 \quad (1\%)$$

Estimation of other marks is roughly as follows:

$$p_{K4me2} = 0.025 \quad (2.5\%)$$

$$p_{K4me1} = 0.10 \quad (10\%)$$

$$p_{K36me3} = 0.17 \quad (17\%)$$

The second question is how to estimate the amount of captured modified nucleosomes. Here we use genomic areas that are not expected to contain marked nucleosomes (based on ChIP-seq of multiple tissues) to estimate the observed background rate $\gamma_{bg}$ in terms of reads/kb (see section above). According to this logic:

$$S_{bg} = \gamma_{bg} * L_{genome} / 1000 \quad \text{and} \quad S_{mod} = S_{total} - S_{bg}$$

Where $S_{total}$ is the total number of unique fragments we recovered and $L_{genome} = 3.3 * 10^9$ is the length of the genome.

Estimating the number of captured fragments from the number of sequenced fragments requires considering the sequencing efficiency --- what fraction of the captured fragments (ones that reached the last PCR step) were actually sequenced. We estimate this efficiency by examining the distribution of duplicate reads in our sequencing output (more below). Given the estimate $\gamma_{seq}$ we estimate

$$C_{mod} = S_{mod} / \gamma_{seq}$$

Putting these all together we get:

$$\gamma_{mod} \approx C_{mod} / N_{mod}$$

$$\gamma_{mod} \approx \left(S_{total} - \gamma_{bg} * 3.3 * 10^6\right) / \left(\gamma_{seq} * p_{mod} * N_{genome} * 1.6 * 10^7\right)$$

And

$$\gamma_{bg} \approx \alpha * 3.3 * 10^6 / \left(\gamma_{seq} * N_{genome} * 1.6 * 10^7\right)$$

The alternative **local method** reasons that every modification has genomic areas which tend to be fully marked and others that are completely unmarked. For instance, nucleosomes flanking promoters of constitutively marked genes (housekeeping genes) are expected to be marked by H3K4me3 in all cells, while areas distant from genes and enhancers are expected to be unmarked. The coverage at these areas depends on the rates aforementioned and thus can be used to estimate cfChIP efficiency.

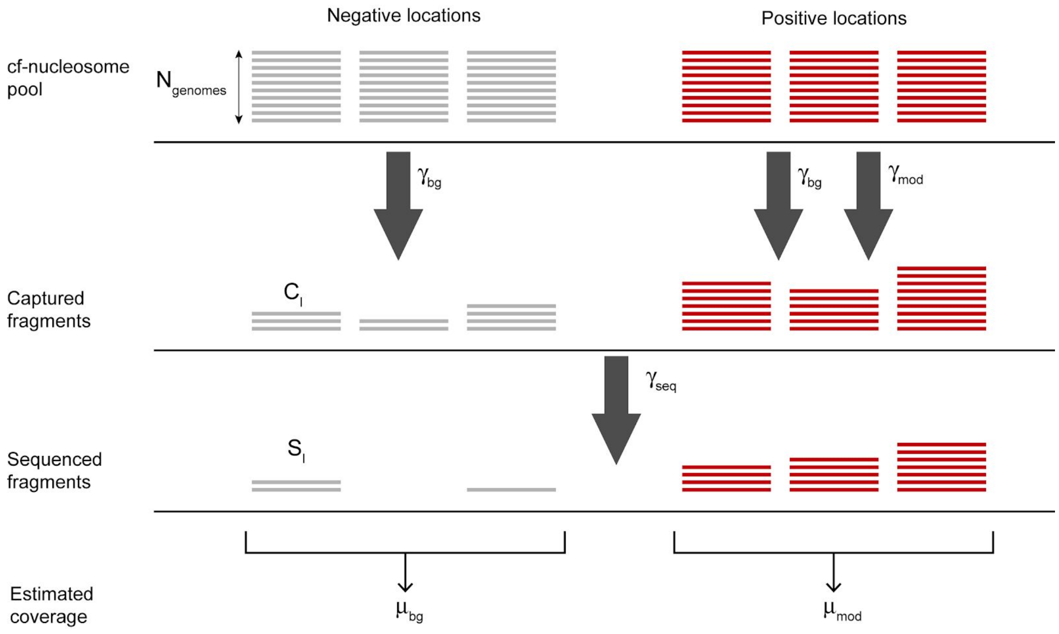


More precisely, let $l$ be a genomic location, and $S_l$ the number of unique sequenced fragments that overlap with it. Then for an unmarked location:

$$E[S_l] = N_{genome} * \gamma_{bg} * \gamma_{seq} \quad \text{(negative location)}$$

We emphasize the expectation, since at each specific location the number of fragments will be distributed around this mean.

In a positive location, where we expect that all input nucleosomes are marked, the expectation is

$$E[S_l] = N_{genome} * (\gamma_{mod} + \gamma_{bg}) * \gamma_{seq} \quad \text{(positive location)}$$

The sum $\gamma_{mod} + \gamma_{bg}$ is due to the fact that a fragment can be captured in a specific or non-specific manner.

To estimate capture rate, we define the coverage in positive location to be the coverage in the 95'th percentile of non background regions

$$Cp = quantile(95, C_{mod})$$

And the coverage in negative locations $\mu_{bg}$ to be the mean coverage at background regions. Using these we conclude that

$$\gamma_{bg} \approx \mu_{bg}/\left(N_{genome} * \gamma_{seq}\right)$$

And

$$\gamma_{mod} \approx Cp/\left(N_{genome} * \gamma_{seq}\right) - \gamma_{bg}$$

Note that this approach circumvents the need to estimate the total number of modified nucleosomes in cells. However, it does require to define positive locations that are assumed to be modified in all cells --- and while there are locations with strong ChIP-seq peaks in multiple cell types that seem to meet that requirement, we do not have any current experimental evidence that support such a claim. However, deviations from the assumption would imply that the estimate is an underestimate of the actual capture rate.

Table S1 contains the estimates using both methods in the samples where we quantified DNA content.

#### Estimation of sequencing efficiency

This problem has been examined in the literature (8) and in various tools (e.g., Picard's EstimateLibraryComplexity). Here we derive a simple method that provides an initial estimate, although more complex ones exist.

We can view the sequencing protocols as starting with a small set of molecules that have the sequencing adapters. These are amplified (16 rounds of PCR) and some fraction of the amplified fragments are sequenced. Due to the large amplification and the numbers of initial fractions (~millions), we can think of the number of times we see a specific input molecule to be Poisson distributed with a parameter $\lambda$ which summarizes the rate at which we see this specific fragment when collecting the actual number of sequences.

In processing the sequenced reads, due to the small amount of input material in cfChIP, we view duplicates as artifacts of the sequencing and not as two identical input molecules. Indeed, comparing two technical repeats from the same plasma sample, we find negligible overlap, suggesting that most duplicates are amplification artifacts.

We can summarize the observed duplicate frequency as

$$\widehat{p}_k = n_k / n_{unique}$$

where $n_k$ are the number of fragments that were duplicated $k$ times, and $n_{unique}$ is the number of unique fragments observed. Estimation of sequencing efficiency can be reduced to estimating $n_0$ the number of fragments we did not observe.

The simplest approach is to assume duplicates are due to a Poisson process with unknown parameter $\lambda$. To estimate this parameter we need a slightly modified procedure, as we do not observe fragments with $k = 0$. This results in a truncated likelihood function

$$l_{poisson}(\lambda) = \sum_{k=1} n_k \log(p(k|\, k > 0,\ \lambda) = \sum_{k=1} n_k \left(\log \tfrac{1}{k!} + k \log \lambda - \log(1 - e^{-\lambda})\right)$$

We can find the maximum likelihood

$$\widehat{\lambda} = \arg\max_{\lambda} l_{poisson}(\lambda)$$

using line search. Once we have this parameter we can estimate the number of total fragments (including the ones we did not see). Briefly, based on the Poisson model,  we conclude that

$$n_{unique} = n_{total} * p(k > 0 \mid \lambda^*) = n_{total} * (1 - e^{-\lambda^*})$$

Where $\lambda^*$ is the unknown real rate. Since sequencing efficiency is the fraction of observed fragments

$$\gamma_{seq} = n_{unique}/n_{total} = 1 - e^{-\lambda^*} \approx 1 - e^{-\widehat{\lambda}}$$

In many cases the actual distribution of duplicates does not match a Poisson distribution. For example

| | |
|---|---|
| 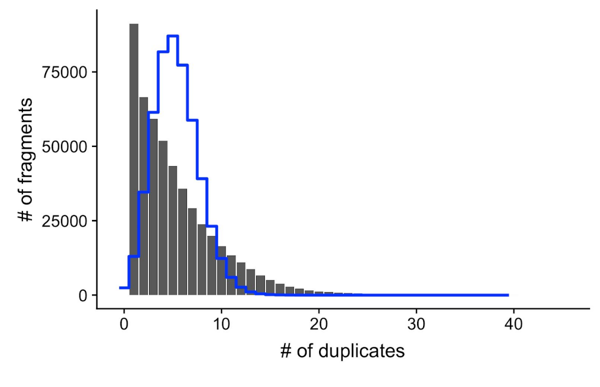  | 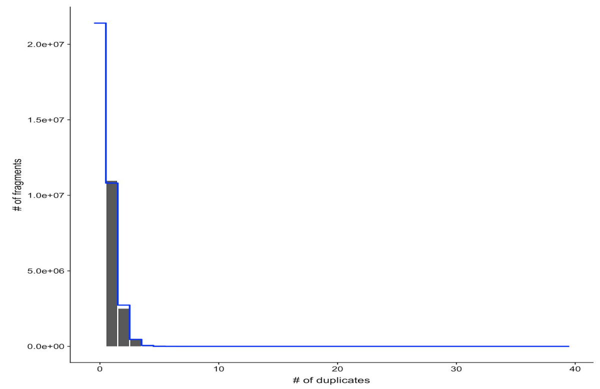  |
| H012.1 - Poisson fit (blue) does not match observed duplication frequency (gray). Note that this sample was sequenced deeply, and the estimate is that there are few unseen | C001.2752 - Poisson fit matches nicely the observed duplication frequency. This sample was sequenced much more shallow and most fragments were seen only once. The estimate |

| | |
|---|---|
| sequences.<br><br>Estimate of $\gamma_{seq}$ is 99.5% | is that there are many more unseen sequences.<br><br>Estimate of $\gamma_{seq}$ is 39.7% |

We reasoned that this discrepancy might be due to some fragments undergoing early duplication(s), thus amplified more than the rest. However, since the actual rate depends on decisions downstream of amplification (e.g., loading libraries onto sequencer etc), we need to assume that the rate of reproducing each copy of these “early winners” is the same as the rest of the fragments. This means that we assume a mixture of Poissons:

$$p_{mix}(k \mid \lambda,\ \rho_1 ..., \rho_M) \ = \ \sum_{m=1}^{M} \rho_m * p_{poisson}(k \mid m * \lambda)$$

Where $\rho_1 ..., \rho_M$ are mixture weights that estimate the proportion of fragments that are amplified once, twice, thrice and so on. Fitting this mixture model requires performing numerical optimization to find parameters that maximize the likelihood. For our two examples we get:

<table>
<tr>
<td>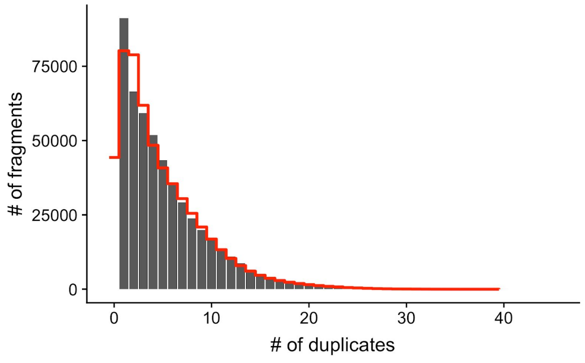
</td>
<td>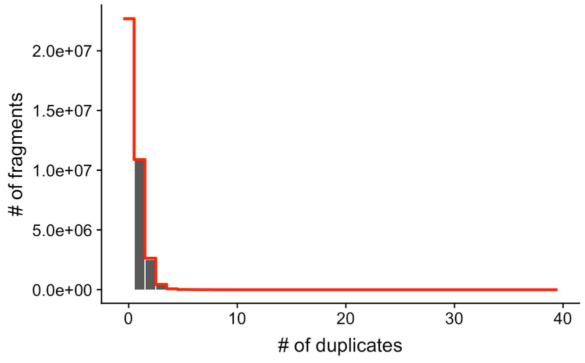
</td>
</tr>
<tr>
<td>H012.1 - with mixture fit (red line). This is much closer to the empirical distribution, and makes a substantially different estimate of the number of unseen fragments.<br><br>Estimate of $\gamma_{seq}$ is 91.4%</td>
<td>C001.2752 - here the mixture fit is close to the Poisson fit above.<br><br>Estimate of $\gamma_{seq}$ is 38.3%</td>
</tr>
</table>

We can gain better sense of these by examining the individual components that make up the mix.

| | |
|---|---|
| 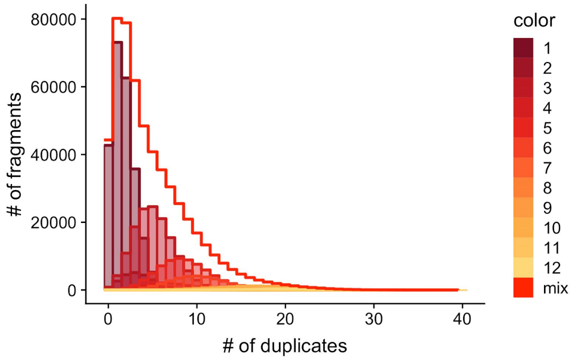  | 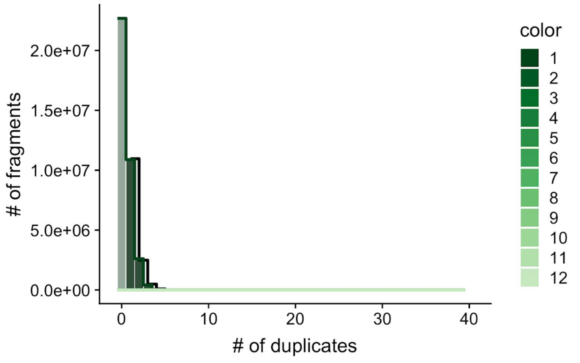  |
| H012.1 - the mixture is made up of several components | C001.2752 - the mixture is dominated by the first component |

Our conclusion (based on these two examples and more) is that for shallow sequenced samples (e.g., similar to C001.2752), the two estimators of efficiency are similar, while for deeply sequenced samples (e.g., similar to H012.1), there are differences in the estimates. Since in these cases the mixture model is closer to the observed duplication frequency we consider it as the more accurate estimate.

To be clear, the estimate of sequencing efficiency according to this model is similar to above:

$$\gamma_{seq} \approx p_{mix}(k > 0 \mid \lambda, \rho_1, ..., \rho_M) = 1 - \sum_{m=1}^{M} \rho_m * e^{-m*\lambda}$$

#### Defining tissue-specific signatures

Using the Roadmap Epigenomics metadata table we defined sets of Roadmap samples that belong to a tissue or group of tissues (see Table S11). These definitions included some redundancies. For example, the group Lymphocytes included B-Cells, T-Cells, and NK samples, and thus subsumed each of these groups.

Recall that $N[w,s]$ is the normalized signal of window $w$ in sample $s$.

We then defined for each group of Roadmap samples the set of specific windows, as windows $w$ meeting the following criteria:

1. The window $w$ is on an autosomal chromosome
2. In at least one of the atlas samples in the group, $N[w,s] \geq 35$
3. In all atlas samples outside the group, $N[w,s] < 15$
4. In all windows $w'$ within 1Kb of $w$, $N[w,s] < 15$

The last condition is added as we noticed that often when a gene is expressed there is “spill over” to neighboring windows.

Groups for which we found less than 4 specific windows were considered to be without signature. For all other groups, we define the signature as the set of specific windows (see Table S4).

#### Statistical tests

Recall that

- $B[w,s]$ - background estimate for window $w$ in sample $s$
- $C[w,s]$ - actual read counts observed in window $w$ in sample $s$
- $\nu[s]$ - normalization factor for sample $s$

**Detection test.** To test whether a gene or a signature is present above background in a sample, we use a Poisson distribution. More specifically:

computeDetectionPValue $(W, s)$

$\lambda \leftarrow \sum_{w \in W} B[w,s]$

$x \leftarrow \sum_{w \in W} C[w,s]$

Return $P_\lambda(X \geq x)$ // Poisson p-value

Here $W$ is a set of windows associated with a gene or a tissue-specific signature as above.

**Estimating mean and variance**. To perform a test of signal vs behavior in a reference set of samples (e.g., samples from healthy cohort), we need to estimate the mean and variance of the signal in the healthy cohort. Ideally, we would want an estimate in units of normalized counts. This estimate is confounded by the sampling noise in each sample which can be a major source of variability, especially for low expression genes. We thus built an estimation procedure that takes this variability into account.

Given a set of windows $W$ of interest, let

- $s_1, \ldots, s_n$ be the set of reference samples
- $c_i \leftarrow \sum_{w \in W} C[w, s_i]$ be the observed count numbers of $W$ in sample $s_i$
- $\lambda_i \leftarrow \sum_{w \in W} B[w, s_i]$ be the estimated background rates of $W$ in sample $s_i$

- $\xi_i = (\nu[s_i])^{-1}$ be the inverse of global normalization factor for sample $s_i$

To estimate the distribution of values, we assume that for each sample

$$C_i \sim Poisson(\xi_i N_i + \lambda_i)$$

Where $N_1, \ldots, N_n \sim P(N)$ are i.i.d. samples of normalized counts from the unknown distribution we want to estimate.

To understand the implications of this model, using linearity of expectations we get

- $E[C_i] = \xi_i E[N_i] + \lambda_i$

This matches how we estimate normalized counts (recall that we defined normalized counts as $N_i \leftarrow (C_i - B_i)/\xi_i$ above).

Using the law of total variation we get that

- $Var[C_i] = \xi_i^2 Var[N_i] + \xi_i E[N_i] + \lambda_i$

which implies that the observed counts variance is due both to the variance of $N_i$ and additional variance due to the observation process.

For the estimate we assume that $N_i$ is distributed according to a Gamma distribution with mean $\mu_N$ and variance $\sigma_N^2$. One often treats the Gamma distribution as the counterpart of normal distributions for non-negative variables. Indeed if $\mu_N \gg \sigma_N$ then the Gamma distribution is almost a normal distribution.

With this assumption, we can use classical results to show that

$$C_i \sim NB(\mu = \xi_i \mu_N,\ \sigma^2 = \xi_i^2 \sigma_N^2 + \xi_i \mu_N) \quad \text{when } \lambda_i = 0$$

Where $NB(\mu, \sigma^2)$ is the Negative Binomial distribution with mean $\mu$ and variance $\sigma^2$. That is,

$$P(Z = k) = \Gamma(k + \tfrac{\mu^2}{\sigma^2 - \mu})\Gamma(k+1)^{-1}\Gamma(\tfrac{\mu^2}{\sigma^2 - \mu})^{-1}(\tfrac{\sigma^2 - \mu}{\sigma^2})^k(\tfrac{\mu}{\sigma^2})^{\mu^2/(\sigma^2 - \mu)} \quad \text{if } Z \sim NB(\mu, \sigma^2)$$

When $\lambda_i > 0$ this is not the precise distribution of $C_i$. However, if $\lambda_i$ is small we can assume that the Negative Binomial is a close approximation to the distribution:

$$C_i \sim NB(\mu = \xi_i \mu_N,\ \sigma^2 = \xi_i^2 \sigma_N^2 + \xi_i \mu_N + \lambda_i)$$

With this assumption we define the likelihood of the parameters we want to estimate

$$L(\mu_N, \sigma_N^2) = \prod_i P_{NB}(c_i\,;\ \mu = \xi_i \mu_N,\ \sigma^2 = \xi_i^2 \sigma_N^2 + \xi_i \mu_N + \lambda_i)$$

We then use an optimization procedure to search for the parameters that maximize the likelihood. (R optim() procedure).

**Comparison of a single sample to a reference cohort test.** Here we test whether the observed signal of a set of windows $W$ in sample $s$ is higher or lower than expected in a reference set of samples (e.g., healthy samples).

We used the following procedure to perform a two-tailed test against the null assumption specified by the given mean and variance.

computePValueAgainstReference( $W$, $s$, $\mu$ , $\sigma^2$ )

$\lambda \leftarrow \sum_{w \in W} B[g,s]$

$x \leftarrow \sum_{w \in W} C[w,s]$

$\mu_X \leftarrow \mu/\nu[s] + \lambda$

$\sigma^2{}_X \leftarrow \sigma^2/\nu[s]^2 + \mu_X$

if( $x > \mu_X$ ) Return $\frac{1}{2}P_{NB(\mu_x,\, \sigma^2{}_X)}(X \geq x)$ // NB p-value

if( $x < \mu_X$ ) Return $\frac{1}{2}P_{NB(\mu_x,\, \sigma^2{}_X)}(X \leq x)$ // NB p-value

As in the test above the set of windows $W$ can be the set associated with a gene or a signature or a group of genes. The difference is that we need to estimate the mean and variance of signal over $W$ in the reference cohort.

**Differential signal between two groups.** Given two groups of samples, $A_1$ and $A_2$ reject the null hypothesis that the signal over a set of windows $W$ is from the same underlying distribution.

We use a *likelihood ratio test* (LRT) where we compare the likelihood under the scenario that all samples in both sets have the same distribution to the alternative where we have a different distribution in each set. The statistics is

$$\chi = 2\left(\log \hat{L}(W,\, A_1 \cup A_2) - \left(\log \hat{L}(W,\, A_1) + \log \hat{L}(W,\, A_2)\right)\right)$$

with

$$\hat{L}(W,\, A) = \max_{\mu,\, \sigma^2} L(\mu_N,\, \sigma_N{}^2 :\, W,\, A)$$

where $L(\mu_N,\, \sigma_N{}^2 :\, W,\, A)$ is the likelihood function defined above with respect to the event defined by $W$ on samples in $A$.

Under the null assumption the distribution of the statistic $\chi$ is approximately a Chi-squared distribution with 2 degrees of freedom, which allows to compute the p-value.

##### Cancer programs

To define cancer signatures from a set of gene programs, we perform the following procedure:

1. For each program $j$ in the set of gene programs, let $W_j$ be the set of windows for the set of genes in the program. Estimate the mean and variance $\mu_j$, $\sigma^2_j$ from the healthy cohort. Then define the p-value of the profile on each sample as

   $p[j,s]$ = computePValueAgainstReference( $W_j$, $s$, $\mu_j$ , $\sigma^2_j$ )

   After computing p-values of all programs against all relevant cancer samples, we define $q[j,s]$ as the q-value of program $j$ in sample $s$ after FDR correction.

2. We define the matrix
   $A[j,s] = 1\{q[j,s] < 10^{-6}\}$
   where $1\{condition\}$ is the indicator function that is equal to 1 when the condition holds and 0 otherwise. We then filter matrix rows (programs) to those that are informative about differences among the samples:
   $A' \leftarrow A\left[\left\{j : 2 < \sum_s A[j,s] < \frac{2}{3}M\right\},\right]$ where $M$ is the number of samples.
3. We cluster the rows of $A'$, and identify for each cluster $k \in \{1, \ldots, K\}$ the consensus over samples as the set of samples where a majority of the programs in the cluster are significant:

   $$A_k = \left\{s : \sum_{j \in Cluster\ k} A'[j,s] > \frac{1}{2}|Cluster\ k|\right\}$$

4. For each cluster $k$ find the set $G_k$ of genes that are differentially marked between $A_k$ and the remaining samples (see above). Use FDR correction to adjust p-values and select genes with $q < 0.001$
5. Repeat steps 1-4 with gene sets $G_1, \ldots, G_K$
6. Return the (revised) gene sets $G_1, \ldots, G_K$ as signatures.
